# Adaptive substitutions underlying cardiac glycoside insensitivity in insects exhibit epistasis *in vivo*

**DOI:** 10.1101/621185

**Authors:** Andrew M. Taverner, Lu Yang, Zackery J. Barile, Becky Lin, Julie Peng, Ana Pinharanda, Arya Rao, Bartholomew P. Roland, Aaron D. Talsma, Daniel Wei, Georg Petschenka, Michael J. Palladino, Peter Andolfatto

## Abstract

Predicting how species will respond to selection pressures requires understanding the factors that constrain their evolution. We use genome engineering of *Drosophila* to investigate constraints on the repeated evolution of unrelated herbivorous insects to toxic cardiac glycosides, which primarily occurs via a small subset of possible functionally-relevant substitutions to Na^+^,K^+^-ATPase. Surprisingly, we find that frequently observed adaptive substitutions at two sites, 111 and 122, are lethal when homozygous and adult heterozygotes exhibit dominant neural dysfunction. We identify a phylogenetically correlated substitution, A119S, that partially ameliorates the deleterious effects of substitutions at 111 and 122. Despite contributing little to cardiac glycoside-insensitivity *in vitro*, A119S, like substitutions at 111 and 122, substantially increases adult survivorship upon cardiac glycoside exposure. Our results demonstrate the importance of epistasis in constraining adaptive paths. Moreover, by revealing distinct effects of substitutions *in vitro* and *in vivo*, our results underscore the importance of evaluating the fitness of adaptive substitutions and their interactions in whole organisms.

## Introduction

Understanding the factors that limit the rate of adaptation is central to our ability to forecast future adaptive evolutionary trajectories and predict the timescales over which these changes are expected to occur (Stern 2011; Losos 2017; Morris et al. 2018). In particular, considerable uncertainty surrounds the relative importance of the availability of adaptive mutations, pleiotropy and epistasis in constraining adaptive paths (Stern 2011; Storz 2018). One fruitful approach to addressing this question has been to examine repeated bouts of adaptation in microbial systems subject to a common selective pressure and identical starting conditions (Jerison and Desai 2015). Unfortunately, such approaches still have limited utility in multicellular eukaryotes and likely don’t reveal the full range of constraints operating in nature. An alternative and analogous approach is to examine evolutionary patterns in large, naturally occurring assemblages of species exhibiting parallel adaptations in response to a common selective pressure (Liu et al. 2010; Meyer et al. 2018; Zhen et al. 2012; Dobler et al. 2012; Christin et al. 2007). Evolutionary studies of parallelisms are a powerful complementary approach to deducing the factors constraining the adaptation (Stern 2013). A well-known example of parallel adaptations is the ability of numerous animals to acquire toxins from their environments and sequester them for use in defense against predators (Brodie 2009; Erb and Robert 2016).

Here, we focus on a large group of herbivorous insects with a broad phylogenetic distribution that have independently specialized on toxic host plants (Dobler et al. 2011). In addition to other defenses against herbivory, the Apocynaceae and other plant species produce a class of toxic secondary compounds called cardiac glycosides (CGs). CGs are highly toxic to animals because they are potent inhibitors of Na^+^,K^+^-ATPase (NKA), a ubiquitously expressed enzyme needed in a variety of cellular processes in animals, including neural signal transduction, muscle contraction, and osmoregulation (Lingrel 2010). Mutations to NKA in invertebrates are typically homozygous lethal and associated with defects in locomotion, neuron development and neural homeostasis (Ashmore et al. 2009). In humans, loss-of-function mutations in NKA have been associated with several rare disorders such as dystonia, parkinsonism and hemiplegic migraines (Bøttger et al. 2012). Despite their toxicity, NKAs have long been targeted with CG-based drugs to treat common conditions such as congestive heart failure and cardiac arrhythmias (Schoner 2002).

Insensitivity to CGs in insects can evolve via several mechanisms including modification of the CG-binding domain of NKA (i.e. target-site insensitivity), restriction of NKA expression to neurons (Petschenka et al. 2013b), the deployment of proteins that ameliorate the toxic effects of CGs (Torrie et al. 2004; Petschenka et al. 2013b) and other physiological factors (Vaughan and Jungreis 1977). Despite this wide variety of potential paths to CG-insensitivity, the evolution of insensitivity in most CG-adapted insects is due, at least in part, to target-site insensitivity. Indeed, in most cases, diet specialization on CG-containing hostplants has been accompanied by recurrent adaptive amino acid substitutions to the CG-binding domain of the alpha-subunit of NKA, ATPα1 (Zhen et al. 2012; Dobler et al. 2012; Yang et al. 2019). Previous studies have identified up to 35 sites in ATPα1 at which substitutions could contribute to CG-insensitivity (reviewed in Zhen et al. 2012). However, CG-insensitivity of ATPα1 most often arises via a highly similar pattern of substitution at two sites (111 and 122, Figure 1A): as an illustration, a survey of 28 CG-adapted insects revealed that 30 of 63 amino acid substitutions observed at sites implicated in CG-sensitivity in ATPα1 occur at sites 111 and 122 (Yang et al. 2019). Sites 111 and 122 have also been identified as targets of positive selection in CG-adapted insects using statistical phylogenetic methods (Yang et al. 2019). Understanding why these two sites, in particular, are so often employed requires a characterization of the effects of these substitutions, individually and in combination, on organismal phenotypes and fitness.

**Figure 1.**
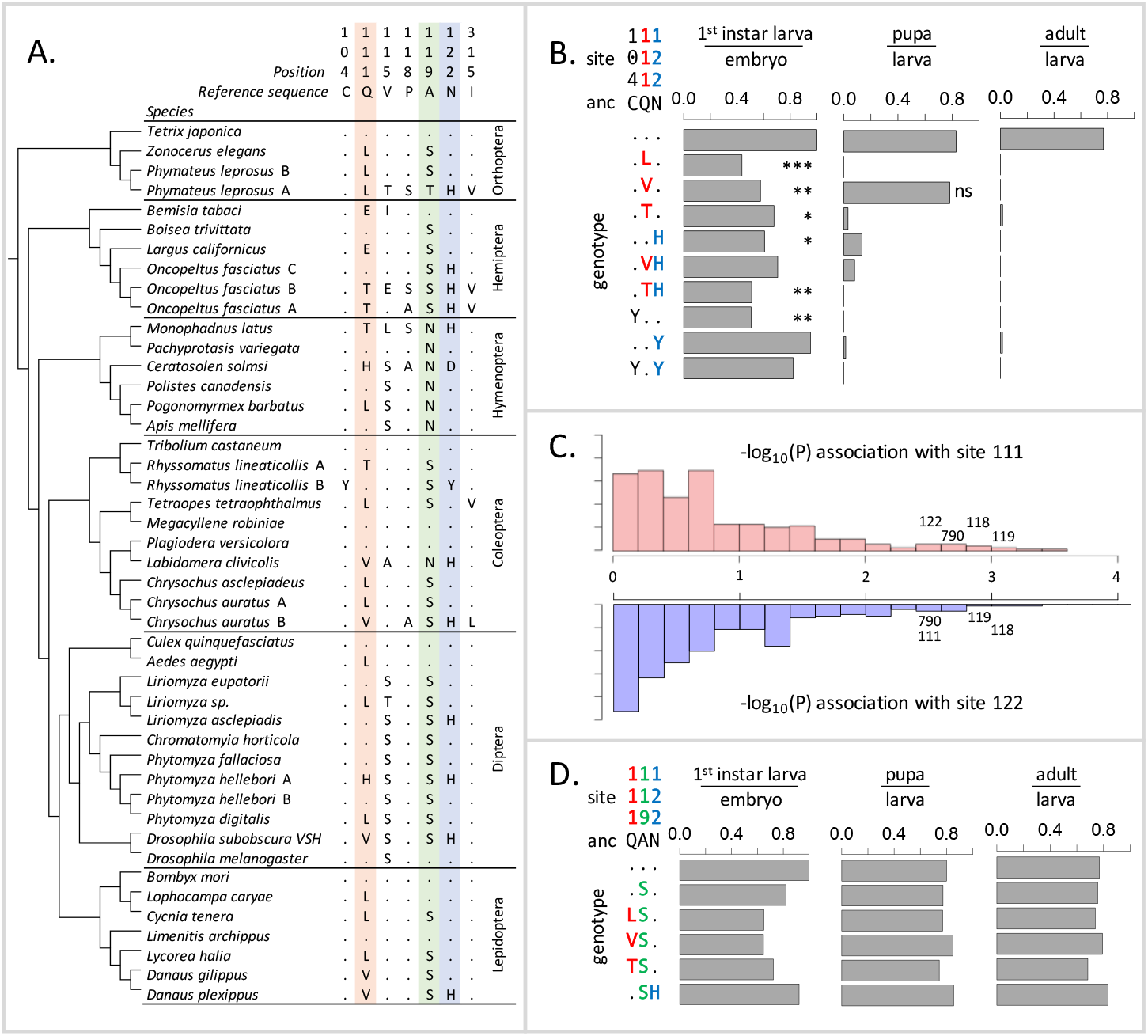
Deleterious effects of substitutions at positions 111 and 122 are ameliorated by the permissive substitution A119S. (**A)** Patterns of amino acid substitution at sites implicated in CG sensitivity near the H1-H2 transmembrane domain of ATPα1 in representative species from six insect orders (see **Figure S2** for all species). Dots indicate identity with the ancestral reference sequence and letters indicate derived amino acid substitutions. Positions 111 and 122, highlighted in pink and blue respectively, are hotspots of frequent parallel substitution. C104Y and N122Y represent rare substitutions associated with ATPα1 duplication (Zhen et al. 2012). Position 119 (highlighted in green) had not been previously implicated in CG sensitivity, but is identified as a candidate permissive substitution in subsequent analyses. (**B)** Viability of homozygous first instar larvae, pupae and adults for the first series of engineered substitution lines. Plotted is the proportion of surviving homozygous mutant offspring (i.e. EYFP-), scaled by the proportion for the wild-type control. Dots indicate identity with the ancestral reference at sites C104, Q111, and N122. Adjusted Fisher’s Exact Test P-values versus the wild-type strain: *** P<0.001; ** P<0.01, *P<0.05. All larvae to pupa survivorship are significantly lower than wild-type (P<1e-5) except for Q111V. All larvae to adult survivorship are significantly lower than wild-type (P<1e-5). (**C)** Distributions of P-values for the strength of the phylogenetic correlation between sites 111 (above) or 122 (below) and 270 non-singleton amino acid variants in a multi-sequence alignment including 174 ATPα1 sequences representing 161 species. Sites 111 and 122 exhibit highly correlated evolution. Three additional sites in the lowest 5% of P-values shared in common between sites 111 and 122, including site 119, are labelled. (**D)** Viability of homozygous first instar larvae, pupae and adults for the second series of engineered substitution lines that include A119S relative to the wild-type control. Dots indicate identity with the ancestral reference at sites Q111, A119, and N122. Despite slight reductions in viability, all reductions are not significant relative to wild-type line after multiple test correction.

To explain the frequent reuse of sites 111 and 122, it has been speculated that substitutions at most alternative sites may be associated with negative pleiotropic effects, that is, have deleterious effects on another aspect of phenotype and fitness (Zhen et al. 2012). Support for this hypothesis comes from the fact that multiple insect species specializing on CG-containing host-plants have independently duplicated and neofunctionalized ATPα1. In all cases examined to date, species with two or more copies retain one minimally altered copy that is more highly expressed in nervous tissue, and have evolved one or more insensitive copies that are more highly expressed in the gut, the site of absorption of CGs (Zhen et al. 2012; Yang et al. 2019). Further support for negative pleiotropic effects is provided by the expression of engineered ATPα1 constructs in cell lines, suggesting that some duplicate-specific CG-insensitivity substitutions appear to reduce NKA activity (Dalla and Dobler 2016). Based on these findings, the frequent parallel substitutions observed at sites 111 and 122 in specialists lacking duplicate ATPα1 copies plausibly reflect the fact that substitutions at these sites are minimally pleiotropic.

### Common substitutions at positions 111 and 122 exhibit negative pleiotropic effects

To test the idea that substitutions at positions 111 and 122 lack strong negative pleiotropic effects, we used the transgenesis toolkit of *Drosophila melanogaster*, a generalist insect that harbors a single ubiquitously expressed copy of a CG-sensitive form of ATPα1. This sensitive form of ATPα1 is the presumptive ancestral state for many potential CG-adapted insects. We focus on several substitutions at sites 111 and 122 (notably Q111V, Q111T, N122H, Figure 1A) have been directly implicated in CG-insensitivity in functional experiments (reviewed in Zhen et al. 2012). By engineering amino acid substitutions into a single *D. melanogaster* background, we ensure that fitness differences observed among lines are caused by the substitution and not confounded by unknown variation in the genomic background. In addition, we can rule out compensatory changes elsewhere in the genome, or evolved changes in physiology, which are concerns in multi-generation population-level experimental evolution studies. Importantly, by testing substitutions *in vivo*, we can evaluate their functional effects at multiple phenotypic levels, from the biochemistry of enzyme inhibition to behavior and fitness. We generated six lines that carry substitutions at these two sites of the endogenous ATPα1 locus individually (Q111L, Q111V, Q111T, N122H) and in combination (Q111V+N122H, Q111T+N122H) (see Methods). For comparison, we also created three lines in which we introduced two rare, copy-specific substitutions (C104Y and N122Y and C104Y+N122Y, Figure 1A) for which we did not have *a priori* expectations about pleiotropic fitness effects. C104Y is known to confer some degree of insensitivity to CGs, and both C104Y and N122Y occur on a neofunctionalised copy of ATPα1 in the milkweed weevil (Zhen et al. 2012). An additional control line carrying the wild-type *D. melanogaster* allele was generated using the same approach.

Based on the repeated use of substitutions at positions 111 and 122, we expected these would confer ATPα1 with some degree of insensitivity to CGs and be associated with either no or mild negative pleiotropic effects on fitness. To our surprise, however, each of the nine lines exhibit severely reduced fitness, behaving effectively as recessive lethals. To identify the developmental stage at which lethality occurs, we evaluated hatchability (i.e. the proportion of homozygous larva per embyro) and the probability of individuals surviving to pupae and adults (Figure 1B). Severe fitness deficits for most engineered substitution lines were apparent as early as the larval hatching stage. Still, for most engineered mutant lines, greater than 50% of individuals hatch into the first instar. Q111V homozygotes exhibit particularly high probabilities of survival until the pupal stage. Nonetheless, survivorship of individuals to adulthood for all lines is close to zero.

### The substitution A119S rescues lethality of substitutions at sites 111 and 122

The unexpected deleterious effects of common substitutions at sites 111 and 122 raises the question of how insects with sensitive ATPα1 isoforms, spanning a broad phylogenetic distribution, can evolve insensitivity to CGs via substitutions to one or both of these sites. An important clue is provided by the observation that *D. subobscura*, a European relative of *D. melanogaster,* carries haplotypes resembling those that we engineered into *D. melanogaster*, including the ancestral (QN) and derived states (QH and VH) at sites 111 and 122, respectively (Pegueroles et al. 2016), and yet are viable. Full length coding sequences for *D. subobscura* (Methods) and the closely related sister species *D. guanche* (Puerma et al. 2018) reveal that 19 amino acid substitutions distinguish the *D. melanogaster* ATPα1 from the ancestral *D. subobscura* protein. We hypothesized that one or more of these substitutions ameliorates the deleterious effects of substitutions at positions 111 and 122.

Using a multiple sequence alignment that includes 161 CG-adapted insects and outgroup species surveyed from six insect orders (Figures S1 and S2, Figure 1A shows a subset of these taxa), we applied the software BayesTraits (Pagel and Meade 2006) to evaluate the evidence for correlated evolution between sites 111 and 122 and all other variant sites in the protein (Methods). Of the divergent sites between *D. melanogaster* and *D. subobscura*, only one site (119) is among those having the strongest phylogenetic correlations with both sites 111 and 122 (top 5% of 270 sites) (Figure 1C, Figure S3). A substitution at 119 is observed in 90% of the cases in which there is a substitution at 111 and 100% of the cases in which there is a substitution at site 122. *D. melanogaster* retains the ancestral Alanine at this site, whereas the *D. subobscura* and closely-related species harbor a derived Serine substitution.

Though site 119 is not among sites known to affect CG-sensitivity (Zhen et al. 2012), previous work identified it as one of four sites in the protein that underlie a CG-association/dissociation rate difference distinguishing human ATP1A1 and ATP1A2 isoforms (Crambert et al. 2004). Considering the orientation of the amino-acid side chain of site 119 with respect to a bound CG (Figure S4), it is unlikely that it plays a direct role in CG binding. Nonetheless, due to its physical proximity to sites 111 and 122 and the evidence for correlated evolution with these sites, we hypothesized that the A119S substitution may compensate for the deleterious effects of substitutions at positions 111 and 122. To test this hypothesis, we generated a second series of substitution lines that include A119S in isolation, and A119S paired with four substitutions at sites 111 and 122 (i.e. Q111L+A119S, Q111V+A119S, Q111T+A119S and A119S+N122H, Figure 1D). Embryos homozygous for A119S, with and without substitutions at 111 and 122, have levels of hatchability and survival that are close to wild-type levels. Remarkably, we find that A119S rescues the lethality associated with homozygosity for all four substitutions with which we paired it (i.e. compare Figure 1D with Figure 1B). These results establish the existence of epistatic fitness interactions between A119S and multiple substitutions at both 111 and 122.

### A119S rescues enzyme dysfunction associated with substitutions at sites 111 and 122

To gain insight into the functional basis of the fitness interaction between A119S and substitutions at sites 111 and 122 at the level of NKA function, we carried out a series enzyme inhibition assays (Figure 2, Figure S5). For lines that are heterozygous for CG-insensitivity substitutions, Mut/+, we expect to see biphasic inhibition curves reflecting the equilibrium dissociation constants (K_d_) corresponding to the wild-type (+) and mutant (Mut) forms of the enzyme, respectively. Comparing biphasic curves for heterozygotes allows us to directly compare inhibition profiles for homozygous-inviable substitutions at positions at 111 and 122 alone and in combination with A119S. Figure 2A details the analysis of the A119S and N122H substitutions in this context. The inhibition curve for A119S/+ appears to be monophasic suggesting that A119S alone has little effect on CG inhibition of NKA. In contrast, the inhibition curve for N122H/+ heterozygotes is biphasic. However, while this implies that N122H substantially increases CG-insensitivity (IC_50,2_ = 6.6e-6), the contribution of the N122H form to the total CG-inhibitable activity in heterozygotes is estimated to be less than half that of the wild-type form (*f* = 0.18, 95% CI 0.14 - 0.24). An analysis of allele-specific expression of N122H/+ indicates close to equal mRNA levels for the two alleles (Figure S6), suggesting that the differences in activity are not due to differences at the level of gene expression. Thus, despite conferring CG-insensitivity, the N122H substitution likely encodes a functionally-impaired enzyme.

**Figure 2.**
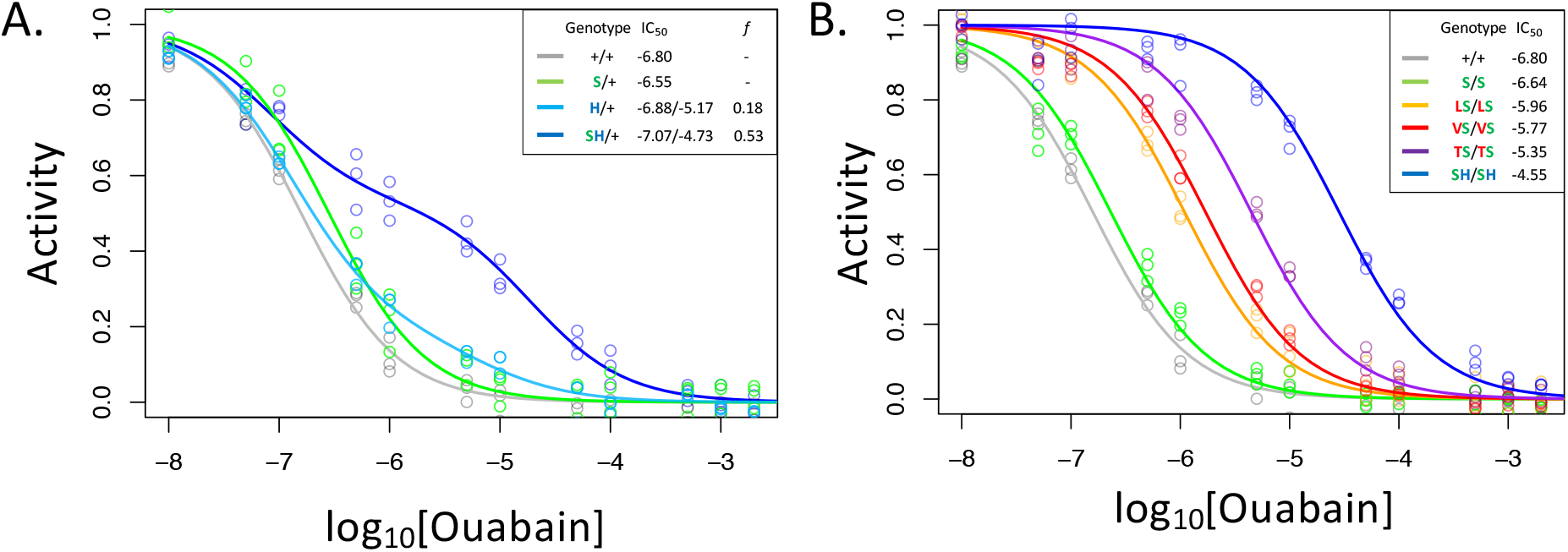
A119S ameliorates detrimental effects of CG-insensitivity substitutions on NKA function. Plotted are relative activity as a function of increasing concentrations of the cardiac glycoside (CG) ouabain. (A) Inhibition curves are plotted for heterozygous individuals to allow comparison of the effects of these substitutions in the presence and absence of A119S. Points represent biological replicates, each of which is the mean across three technical replicates. Curves for the engineered wild-type strain (+/+) and A119S/+ (S/+) are plotted for comparison. The presence of A119S alone confers a negligible increase in CG-insensitivity and results in a monophasic curve. In contrast, N122H/+ (H/+) and A119S+N122H (SH/+) exhibit a biphasic curves. The estimated proportion of CG-inhibitable activity of the mutant form for H/+, *f* = 0.18 (95% CI 0.14 - 0.24), is significantly lower than that of SH/+, *f* = 0.53 (95% CI 0.47 - 0.56). (B) Inhibition curves for homozygous substitutions Q111L, Q111V, Q111T and N122H in the presence of A119S reveal that they increase CG-insensitivity by 7, 11, 28 and 178-fold, respectively. The effect of A119S alone is estimated to be 1.45-fold relative to +/+ (95% CI 1.13 – 1.86).

In contrast to N122H/+, N122H+A119S/+ heterozygotes produce a strongly biphasic inhibition curve, with a two-fold higher estimated IC_50,2_ = 1.9e-5 and comparable levels of activity to that of the wild-type form (*f* =0.53, 95% CI 0.47-0.56, Figure 2A). Similar results were obtained in comparisons of Q111V and Q111V+A119S (Figure S5). We estimate the level of CG-insensitivity conferred by Q111L, Q111V, Q111T and N122H, in the presence of A119S, to be 7, 11, 28 and 178-fold greater than the wild-type form of the enzyme, respectively (Figure 2B). Our results demonstrate epistasis between substitutions at sites 119 and 111/122 at the level of enzyme function. Previous studies have demonstrated increased survival of cell lines expressing Q111V, N122H, and Q111T+Q111H (Dobler et al. 2012), and substantial insensitivity to CG-inhibition despite little effect on ATPase activity for N122H, Q111T+Q111H and Q111V+Q111H (Dalla et al. 2013; Dalla and Dobler 2016). Given that these substitutions were engineered on a *D. melanogaster* background lacking A119S, our results suggest such substitutions are likely to be associated with substantial enzyme dysfunction.

### A119S rescues neural dysfunction associated with substitutions at sites 111 and 112 *in vivo*

An advantage of functionally testing the effects of amino acid substitutions *in vivo*, as opposed to *in vitro*, is the ability to examine fitness-related phenotypes at multiple levels. We therefore considered the impact of substitutions, alone and in combination, on higher level fitness-related phenotypes including adult behavior in response to stress and survivorship of adult flies exposed to CGs.

NKA plays a central role in maintaining neuron action potentials and previous studies have documented short-term paralysis following mechanical overstimulation (aka “bang sensitivity”) associated with mutations that reduce this enzyme’s activity (Ganetzky and Wu 1982; Schubiger et al. 1994; Davis et al. 1995; Palladino et al. 2003). As such, we used the bang sensitivity phenotype as a proxy for proper neural function. Examining our panel of substitutions for these effects reveals all of the substitutions we engineered at sites 111 and 122, individually and in combination, exhibit dominant bang sensitivity phenotypes that are similar in severity to those of the loss-of-function deletion Δ2-6b (Figure 3A, Figure S7). In contrast, bang sensitivity phenotypes were indistinguishable from wild-type for all heterozygous substitutions at 111 and 122 in combination with A119S (Figure 3A, Figure S8). Interestingly, individuals homozygous for A119S, and substitutions at 111 and 122 in the presence of A119S, still exhibit obvious neural dysfunction relative to the wildtype (Figure 3, Figure S8). Thus, while A119S is itself associated with recessive pleiotropic effects, our results demonstrate some degree of positive epistasis between A119S and substitutions at sites 111 and 122 when considering the level of adult behavior.

**Figure 3.**
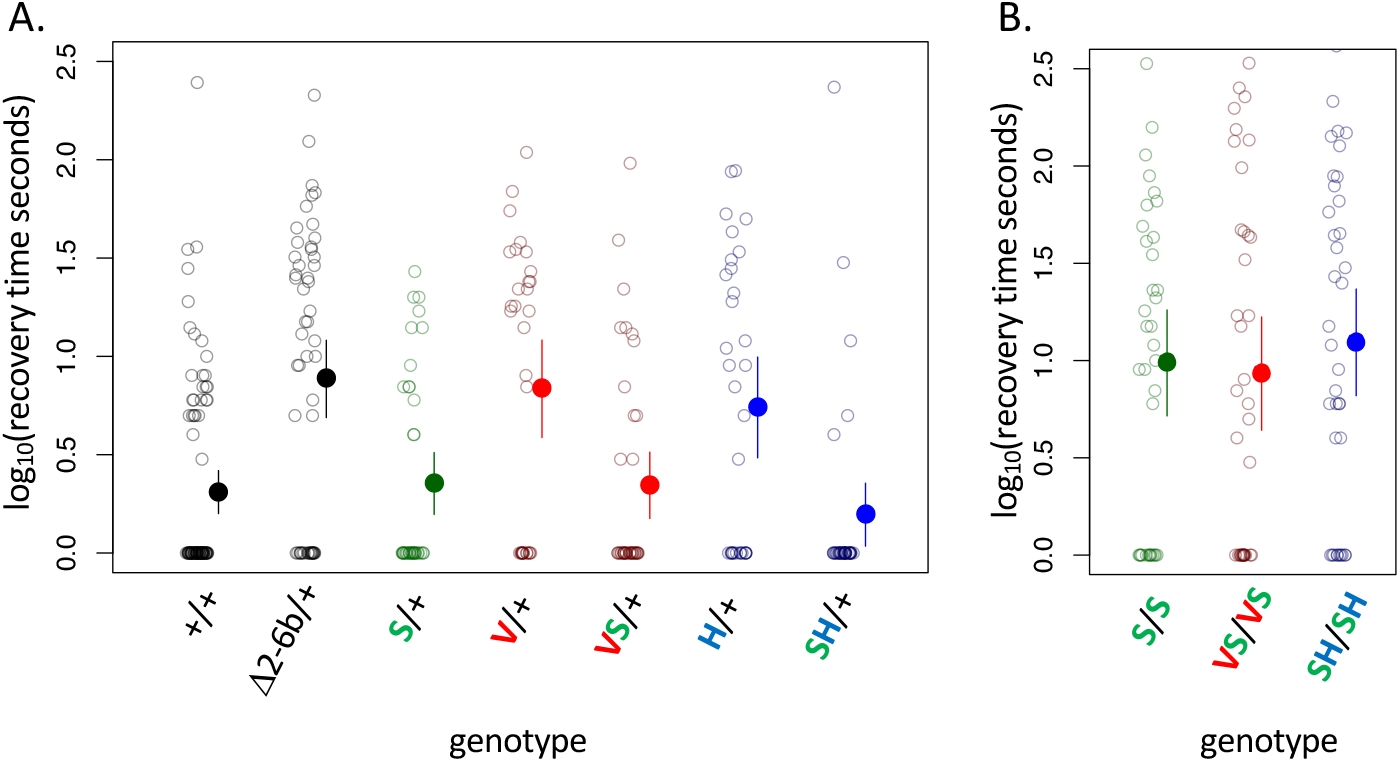
A119S ameliorates detrimental effects of CG-insensitivity substitutions on neural function. Plotted are recovery times for individuals (open circles), and means with approximate 95% confidence bounds (solid circles with whiskers), following mechanical over-stimulation (aka, the “bang sensitivity” assay). Flies heterozygous for loss-of-function ATPα1 substitutions are known to have significantly longer recovery times than wild-type flies (see for e.g. Δ2-6b/+ relative to +/+ in panel A, *P*=4.5e-7). **(A)** Distributions of recovery times for heterozygous flies (i.e. Mut/+) carrying individual substitutions A119S (S/+), Q111V (V/+) and N122H (H/+) and combinations VS/+ and SH/+. The recovery-time distributions for S/+, VS/+ and SH/+ are indistinguishable from the engineered wild-type strain (+/+). In contrast, recovery times are significantly longer than wild-type for lines with individual substitutions V/+ and H/+ (*P*=1.3e-5 and *P*=5.2e-4, respectively) and are indistinguishable from the loss-of-function deletion mutation Δ2-6b/+. (B) Recovery times for homozygous lines S/S, VS/VS and SH/SH are all significantly longer than for heterozygotes S/+, VS/+, and SH/+ (*P*<3.3e-3). These results demonstrate epistasis between substitutions at sites 119 and 111/122 at the level of neural function. A119 ameliorates the dominant dysfunction caused by Q111V and N122H but residual recessive neural dysfunction is still apparent in homozygotes that include A119S.

### Substitutions at sites 111, 119 and 122 increase adult survival upon exposure to CGs

*D. melanogaster* do not normally consume CG-containing plants and consumption of CGs results in increased mortality (Groen et al. 2017). Given that substitutions at sites 111 and 122 decrease sensitivity to CG-inhibition of NKA, such substitutions should confer an advantage upon exposure of *D. melanogaster* to CGs. To test this hypothesis, we exposed adult animals to media containing the CG ouabain. While the wild-type strain suffers high levels of mortality upon CG exposure, the lines carrying substitutions at sites 111 and 122 in combination with A119S are all substantially less-sensitive (Figure 4, Figure S9). Notably, the survival probability of A119S+N122H (SH/+ and SH/SH) is indistinguishable from control treatments in exposures with up to 10 mM ouabain. Beyond confirming an association between insensitivity to CG-inhibition of enzyme activity *in vitro* and reduced sensitivity to CG-exposure *in vivo*, two additional important findings arise from these experiments. First, a substantial measure of insensitivity to CG-exposure is also conferred by A119S alone (S/+ and S/S exhibit ∼8.5-fold and ∼21-fold lower relative risk, respectively, than wild-type at 5 mM ouabain, Figure 4). Despite this, A119S alone has only a small effect on NKA sensitivity to CG-inhibition (Figure 2). Further, the substantially improved survival of heterozygous strains relative to the wild-type strain, including to some extent A119S/+, suggests that insensitivity to CG exposure is a partially dominant phenotype (Figure 4, Figure S9). Importantly, despite the lethal homozygous effects of Q111V and N122H individually (Figure 1B), the dominant effects of these substitutions on insensitivity to CG-exposure imply that they have the potential to confer a fitness advantage as heterozygotes in CG-rich environments.

**Figure 4.**
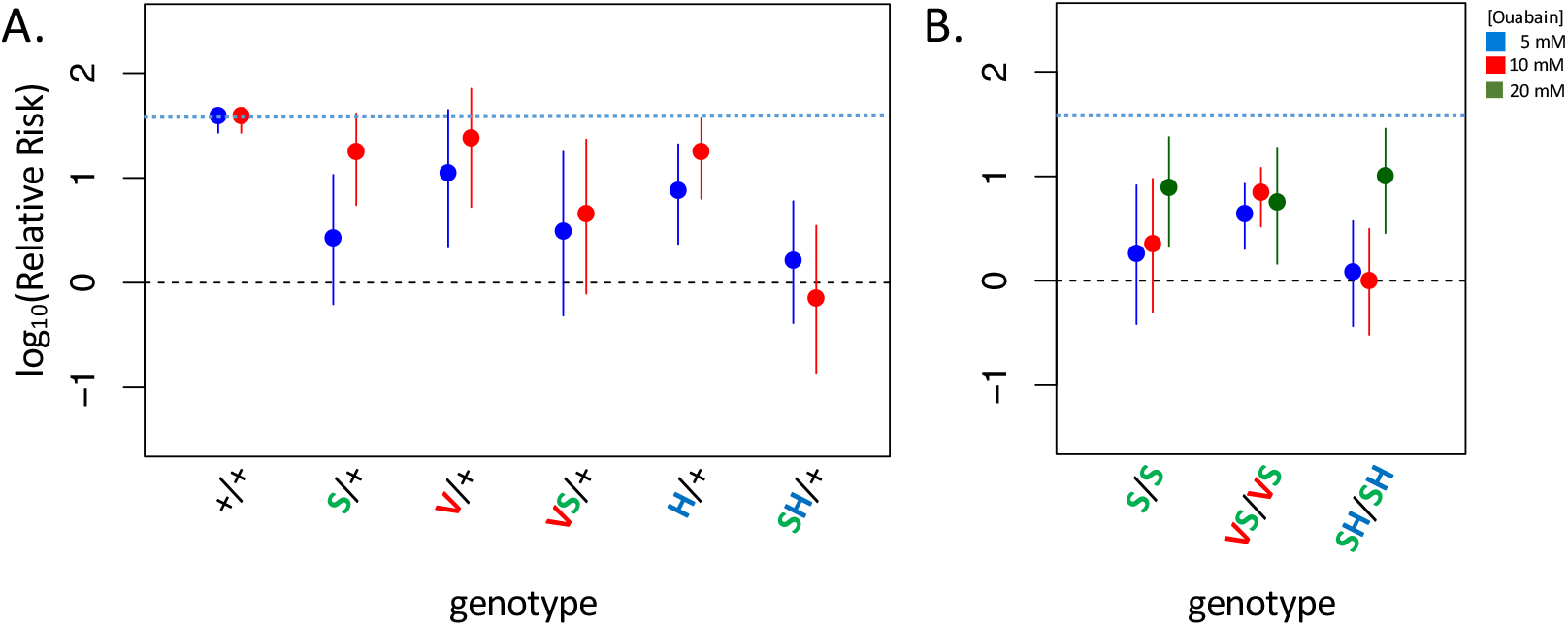
Adult survival upon 7-day exposure to the CG ouabain. Plotted is the log relative risk for treatments (5, 10 or 20 mM ouabain) relative to no treatment controls (no ouabain) for (A) heterozygous strains and (B) homozygous strains. Estimates (points) and 95% confidence bounds (whiskers) were obtained using the Cochran-Mantel-Haenzel framework. Each estimate is based on three biological replicates of 20 flies per concentration. A value of 0 corresponds to equal probability of survival on treatment versus the no ouabain control indicating complete insensitivity to a given concentration of ouabain. Labels: + = engineered wild-type allele; S = A119S; V = Q111V; VS = Q111V+A119S; H = N122H; SH = A119S+N122H. These results reveal appreciable CG-insensitivity conferred by individual substitutions A119S (S/+ and S/S) and the recessive lethals Q111V (V/+) and N122H (H/+) compared to the engineered wild-type control strain (+/+).

### Implications for the evolution of CG-insensitivity in insects

Our findings carry important implications for how CG-insensitivity evolves in insects. At the onset of this study, we expected minimal negative pleiotropic effects associated with adaptive substitutions at sites 111 and 122, given their frequent parallel occurrence across insect orders (Zhen et al. 2012). Our findings that these substitutions are associated with negative pleiotropic effects at multiple levels (i.e. enzyme function, neural function and viability) reveals a more complicated explanation. As we show, with few exceptions, adaptive substitutions conferring CG-insensitivity at sites 111 and 122 most often arise on a genetic background that includes a substitution at site 119, a site not previously implicated in CG-sensitivity. We further show that the most commonly observed substitution at sites 119, A119S, rescues the lethality associated with substitutions at sites 111 and 122, as well as their dominant effects on neural function. A second common substitution, A119N, appears independently in three insect orders and may act similarly (Figure 1A, Figure S2). Thus, the repeated use of specific substitutions does not mean that these changes unconditionally lack negative pleiotropic consequences and that their fitness advantage can depend critically on the background on which they arise.

To the extent that *D. melanogaster* represents a typical CG-sensitive species, it is clear that substitutions at positions 111 and 122 would not be permitted to fix in the species without being preceded by A119S or an equivalent permissive substitution. A119S was not detected in extensive mutagenesis screens (Zhen et al. 2012) and has only modest effects on CG-inhibition of NKA *in vitro* (Figure 2). Given these findings and its position and orientation in the NKA-ouabain co-crystal structure (Figure S4), one might assume that A119S is a neutral (or nearly-neutral) permissive substitution that renders some species candidates for the adaptation to CGs by chance. However, our ability to examine the effects of A119S, and interacting substitutions, in the context of whole animal phenotypes reveals a remarkably different picture. In particular, we show that homozygosity for A119S results in substantial levels of neural dysfunction (Figure 3). Despite these detrimental effects, we also find that A119S, even when heterozygous, confers a substantial survival advantage upon exposure to CGs. Such insights could only be provided by evaluating the effects of mutations in the context of the whole organism. Investigating just one aspect of phenotype, for example biochemistry *in vitro*, yielded a misleading picture of the fitness advantages, and disadvantages, that likely operate on the individual amino acid substitutions underlying this adaptation.

Our findings raise the question of how, given its deleterious effects, A119S became established in so many species. Initially, a survival advantage conferred by A119S in CG-rich environments may have outweighed deleterious effects associated with homozygosity for this substitution. This may be especially true for insects that specialized on CG-containing plants, where competition with other species for resources is reduced. In multiple lineages, the substitutions A119S and A119N preceded substitutions to 111 and 122 (Hemiptera, Hymenoptera, Diptera). In Drosophila, where we have the greatest phylogenetic resolution, A119S was established before substitutions to sites 111 and 122 in the evolutionary lineage leading to *D. subobscura*. This said, Q111L is sometimes observed in the absence of A119S (as is Q111E, see Figure 1A: *Bemisa tabaci* and *Lophocarya caryae;* not shown are *Aedes aegypti*, *Mechanitis polymia* and *Empyreuma pugione*, see Figure S2) implying alternative permissive substitutions in these taxa. More generally, A119S is not a particularly uncommon substitution among taxa that do not specialize on CG-containing hostplants (Figure S2), again implying the existence of additional compensatory substitutions in these species. None of the 18 sites distinguishing the *D. subobscura* and *D. melanogaster* proteins appear among sites most highly correlated with A119S in a phylogenetic analysis, however, leaving us few clues about promising candidates (Figure S3).

This said, a model invoking serial substitution of A119S and compensatory mutations is not the only plausible scenario. Alternatively, since CG-insensitive haplotypes substitutions including A119S lack strong deleterious effects and may confer a survival advantage upon exposure to CGs when heterozygous (Figure 3, Figure 4, Figure S8), such haplotypes may be maintained over time by heterozygote advantage and subsequently reach fixation as they accumulate a sufficient number of compensatory substitutions (Kimura 1985) or by gene duplication (Spofford 1969), which has occurred repeatedly in CG-specialist species (Zhen et al. 2012; Petschenka et al. 2017; Yang et al. 2019). Interestingly, *D. subobscura* is polymorphic for substitutions at positons 111 and 122 mediating CG-insensitivity and these haplotypes are associated with polymorphic inversions that may rarely be homozygous (Pegueroles et al. 2016). The evolution of CG-insensitivity in *D. subobscura* may be recent and represent a transient stage in the evolution of this adaptive trait.

Our study adds to a growing body of work linking epistasis to the dynamics of adaptive evolution in contexts where the ecological relevance of the phenotype is well-understood (Tarvin et al. 2017; McGlothlin et al. 2016; Tufts et al. 2015; Kumar et al. 2017). To date, however, epistasis among substitutions in a protein has been evaluated *in silico*, *in vitro*, and in microbes along the dimensions of either protein folding stability, ligand specificity, enzyme activity, or cell growth rates (Storz 2016). While it is tempting to use one axis of the mutation-phenotype map as a convenient proxy for fitness (e.g. *in vitro* studies of the effects of mutations on protein function), the actual effect of a given mutation on fitness is a convolution over multiple phenotypes, from the levels of enzyme and tissue function to that of whole organism fitness in its environment. By considering phenotypic effects both *in vitro* and *in vivo*, we demonstrate how dominance and epistasis of amino acid substitutions can be missed or mischaracterized in the absence of a whole animal model to evaluate these effects. As such, our study reveals surprising new features of adaptive amino acid substitutions that were previously inaccessible.

## Methods

### Sequencing and alignments

Sources of data used in this study are detailed in Table S1. We collected new data for three species: *Drosophila subobscura* (Diptera), *Monophadnus latus* (Hymentoptera), and *Syntomeida epilais* (Lepidoptera). In each case, total RNA was extracted using TRIzol (Ambion, Life Technologies) following the manufacturer’s protocol. RNA-seq libraries were prepared with either the TruSeq RNA Library Prep Kit v.2 (Illumina) or TruSeq Stranded RNA Library Prep Gold (Illumina) and sequenced on HiSeq2500 (Genomics Core Facility, Princeton, NJ, USA). 50-55 million paired-end 150 nucleotide reads per library were trimmed for quality (Phred quality ≥ 20) and length (≥30 contiguous bases). Trinity 2.2.0 (Haas et al. 2013) was used for *de novo* transcriptome assembly using default parameters. The ATPα1 coding sequences of *D. melanogaster* (FBgn0002921) and *B. mori* (GenBank: LC029030.1) were used to query the assembled transcripts using either tblastx or tblastn (blast-2.26). For *Dysdercus cingulatus* (Hemiptera), *Macrocheraia grandis* (Hemiptera)*, Largus californicus* (Hemiptera), *Jalysus sp.* (Hemiptera), *Metatropis rufescens*, *Ischnodemus falicus*, *Geocoris sp.*, *Gyndes sp.*, *Monomorium pharaonis*, *Hylobius abietis*, *Megacyllene caryae*, *Calligrapha philadelphica*, and *Basilepta melanopus,* we downloaded raw RNA-seq data, carried out *de novo* assembly and obtained ATPα1 sequences using methods similar to those described above. The remaining ATPα1 sequences were either previously published or obtained from publicly available genome references assemblies. A multiple nucleotide sequence alignment was created using ClustalOmega (Sievers et al. 2011). Two alternatively spliced exons were masked for subsequent phylogenetic analyses.

### Statistical phylogenetic analyses

Phylogenetic relationships were established based on previously published sources (Table S2). Phylogeny branch lengths were estimated using IQtree (Nguyen et al. 2015) on the nucleotide alignment of 174 ATPα1 sequences representing 161 species sampled from 8 insect orders (Figure S2) using a guide phylogeny (Figures S1) to force species branching order within orders. We carried out a comprehensive search for substitutions at ATPα1 on which substitutions at sites 111 and 122 were dependent using BayesTraits (Pagel and Meade 2006). We first reconstructed the ancestral sequences of ATPα1 using PAML (Yang 2007). As inputs, we provided PAML with the phylogeny with branch lengths and the amino acid alignment for ATPα1. We used the default PAML parameters with the following changes:“cleandata” was set to 0 to preserve sites with missing data; “fix_blength” was set to 1 to use the branch lengths as the initial values in the ancestral reconstruction; “method” was set to 1 to use a new algorithm in PAML that updates branch lengths one-by-one. Using the inferred ancestor, we then binarized each amino acid state in the multi-species alignment into ancestral, “0”, and derived, “1”, and used this, and the phylogeny with branch lengths as the input for BayesTraits. BayesTraits fits a continuous-time Markov model to estimate transition rates between discrete, binary traits and calculates the likelihood of the fitted model. Restricting rate parameters appropriately (see below), we tested whether the transition rates for sites 111 and 122 were dependent on the state at all other variant sites. We excluded 56 sites with just a single substitution (i.e. one instance in 174 sequences in the alignment) due to their low information content.

In all models, double transition rates (e.g. Q111V and A119S occurring at the same time) were set to zero as double transitions are highly unlikely to occur in a single step and can be modeled as two single transitions, following Pagel (1994). Additionally, we set all transition rates from the derived state back to the ancestral state to zero as failing to do this resulted in unrealistically high estimates of the reversion rate. After these restrictions were imposed, the null model (i.e. independence) had four transition parameters. To test the dependence between sites, we refit an alternative model with two additional restrictions: one forcing the transition rate at site 1 to be fixed regardless of the state of site 2, and a second forcing the transition rate at site 2 to be fixed regardless of the state of site 1. This effectively tests whether the transition rate is affected by the state of either site.

Following the BayesTraits manual recommendations, the phylogeny branch lengths were scaled using BayesTraits to have a mean length of 0.1. Additionally, to increase the chance of finding the true maximum likelihood, we set MLTries to 250 which controls the number of times BayesTraits calls the maximum likelihood algorithm, returning the maximum likelihood of the 250 attempts. We ran each pair 25 times and checked that at least half of the runs had equivalent maximum likelihoods (within 0.1) to ensure that the results were stable. Taking the maximum likelihood for each pair, for each model, we calculated p-values using the LRT, where the statistic of merit is: 2 (unrestricted model – restricted model). The LRT statistic is distributed chi-squared with degrees of freedom (df) equal to the number of restrictions imposed on the model (df=2).

### Engineering ATPα amino-acid substitution lines

To test for phenotypic effects associated with candidate amino-acid substitutions, we developed a genetic engineering approach of the endogenous *ATPα1* (CG5670) locus in *Drosophila melanogaster*. Employing a strategy similar to that of Roland et al. (2013), we first generated a “founder line” by deletion of exons 2-6b and replacement with a functional *attP* element using ends-out homologous recombination (Figure S10). To generate the founder line, homology arms were generated as 2-3 kilobase PCR amplicons 5’ of exon 2 and 3’ of exon 6b and inserted into the *pGX-attP* vector (Huang et al. 2009). This construct was injected into *w^1118^* embryos using standard protocols for P-element transgenesis (Rainbow Transgenic Flies, Inc.). Ends out recombination was performed and molecularly confirmed. The *mini-white* gene was subsequently removed by Cre-lox mediated excision (Groth et al. 2004) to create the founder line, *w^1118^;;ATPαΔ2-6b attP/TM6B,Tb1* (exons 2-6b were replaced by a functional *attP* site).

To generate allelic variant lines of ATPα1, we cloned exons 2-6b into *pGX-attB-ATPα2-6b* vector and injected it into the *attP* founder line mated to *vasa-phiC31-ZH2A* flies using standard protocols (Rainbow Transgenic Flies, Inc). The unmodified plasmid was to generate a wild-type control (*GE-ATPα-D2-6b-WT*) following Cre-lox reduction of the *mini-white*. We then used site-directed mutagenesis (Quick-change Lightening, Agilent) to generate a panel of amino-acid substitution variants of *pGX-attB-ATPα2-6b* and corresponding GE strains were generated and Cre-lox reduced, as above for the WT control (Table S3). Variant and control lines were balanced by crossing to *w;;Dr*[*1*]*/TM6B, P{w*[*+mC*]*=Dfd-EYFP}3, Sb^1^,Tb^1^,ca^1^* (Bloomington Stock Center line 8704) and selecting for EYFP florescence. Homozygous variant GE-ATPα lines were established, when possible, by sib-mating and selection of non-florescent *Tb^+^,Sb^+^* larvae. All substitutions were confirmed using PCR and Sanger-based sequencing.

### Viability assays

Viability was measured in three ways. First, we measured viability as the relative hatchability of embryos carrying a given homozygous amino acid substitution (hereafter “Mut”). To do this, we self-crossed each balanced *GE-ATPα-D2-6b-Mut/TM6B, P{w*[*+mC*]*=Dfd-EYFP}3, Sb^1^,Tb^1^,ca^1^* and allowed them to lay fertilized eggs on standard apple juice–agar Petri plates with yeast paste for two hours at 25°C, 50% humidity. After 24 hours, the relative proportion of non-EYFP (i.e. homozygous *Mut*/*Mut*) to EYFP (i.e. heterozygous *Mut*/+) first instar larvae were counted. As a control, we generated *GE-ATPα-D2-6b-WT/TM6B, P{w*[*+mC*]*=Dfd-EYFP}3, Sb^1^,Tb^1^ parents* and followed the same procedure. A second measure of viability is the probability of survival of a first instar *Mut*/*Mut* larvae to adulthood. To measure this, we transferred homozygous (*Mut/Mut*) first instar larvae to fly media vials (recipe R, LabExpress, Ann Arbor, MI). Larval density was limited to 10 per vial and vials were kept at 25°C, 50% humidity. After two weeks, we counted the number of emerging adults. Homozygous *GE-ATPα-D2-6b-WT* first instar larvae generated the same way were used as a control. Third, we measured egg-to-adult fitness as the proportion of emerging *Mut/Mut* adults in media bottles (recipe B, LabExpress, Ann Arbor, MI) seeded with *GE-ATPα-D2-6b-Mut/TM6B, P{w*[*+mC*]*=Dfd-EYFP}3, Sb^1^,Tb^1^,ca^1^* parents and used bottles seeded with *+/TM6B, P{w*[*+mC*]*=Dfd-EYFP}3, Sb^1^,Tb^1^,ca^1^* parents as controls.

Homozygous mutant (i.e. EYFP-) embryos represent ∼1/4 of screened embryos. For series 1 engineered lines, Figure 1B, mean sample sizes for EYFP-individuals were 243 (range 153-344) for 1^st^ instar larva | embryo, 62 (range 41-117) for pupa|larva, and 57 (range 31-117) for adult | larva. For series 2 engineered lines, Figure 1D, Mean sample sizes for EYFP-individuals were 203 (range 126-478) for 1^st^ instar larva | embryo, 153 (range 99-344) for pupa|larva and adult | larva.

### Enzyme inhibition assays

For each *D. melanogaster* substitution line, we homogenized 90 heads (previously stored at −80°C) in 900 µl of deionized water, using a 1 ml glass grinder (Wheaton) chilled on ice (Petschenka et al. 2017). After vortexing, we divided homogenates into three aliquots of 300 µl representing technical replicates. Subsequently, samples were frozen at −80°C and freeze-dried (Christ, Alpha 2-4 LDPlus) overnight. For assessing resistance of NKA to ouabain, we followed the procedure as described in (Petschenka et al. 2013a) that is based on the photometric evaluation of phosphate released from enzymatic hydrolysis of adenosine triphosphate (ATP) as a measure of NKA activity. Lyophilisates were stored at −80°C until use and reconstituted with 500 µl deionized water. Head extractions were incubated at increasing concentrations of the water-soluble standard cardenolide ouabain (10^−8^ M to 2 × 10^−3^ M, Sigma, Germany, O3125-1G) at 37°C under the following conditions: 100 mM NaCl, 20 mM KCl, 4 mM MgCl_2_, 50 mM imidazol, and 2.5 mM ATP (pH 7.4). We corrected all measurements for a background value that we obtained by incubating the extract under the same conditions as above, except with the addition of 2 × 10^−3^ M ouabain and no KCl (i.e., NKA inactive). On each microplate, we included an assay with porcine Na^+^/K^+^-ATPase (Sigma, Germany, A7510-5UN) as an internal standard. In addition, we ran a series of KH_2_PO_4_ dilutions as a phosphate calibration curve on every plate. We measured absorbances at 700 nm using a CLARIOstar microplate reader (BMG Labtech, Germany). We carried out three biological replicates per line based on different extractions of fly heads. Each biological replicate was the average of three technical replicates, i.e. measurements based on aliquots from original extracts. For data analysis, we compared all measurements to a non-inhibited control. We fitted inhibition curves using OriginPro 2017 (OriginLab, Northampton, MA) with top and bottom asymptotes set to 100 and 0, respectively. Alternatively, we used least-squares fitting (using the nlsLM function of the minpack.lm library in R) to the bi-phasic curve function,

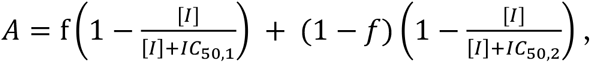

where IC_50,1_ and IC_50,2_ represent the inhibitor concentration [I] required for 50% activity *A* of each isoform present in proportions *f* and *(1-f)*, respectively. Setting *f*=0 assumes a homogenous population of enzyme as expected for homozygotes. The two methods yielded similar IC_50_ estimates. Confidence intervals on estimates of IC_50,1_, IC_50,2_ and *f* were estimated using parametric bootstrap simulations, assuming that residuals across biological replicates are normally distributed.

### Targeted allele-specific expression

To estimate relative expression of mutant and wild-type alleles in heterozygous strains, we designed a targeted allele-specific expression (ASE) assay. Genomic DNA (gDNA) and total RNA were extracted sequentially using TRIzole Reagent (Invitrogen 15596026 and 15596018) from three replicate pools of 20 males sampled from each of sixteen lines. Extracted RNA was further purified with Qiagen RNAeasy column (Qiagen 74104) and reverse transcribed into cDNA with the use of random primers and ProtoScript II Reverse Transcriptase (NEB M0368S). Two sets of primers for ATPα1 were designed and used in subsequent PCR, multiplexing and sequencing. The first set were designed as exon-primed intron crossing (EPIC, Racle et al. 2017) primers and span a short intron separating exons 2 and 3. A second set of primers was used to add standard Illumina-like i5 and i7 barcodes to the PCR amplicons to facilitate multiplexing and subsequent sequencing. These multiplexed amplicons were pooled and sequenced on an Illimina MiSeq (Princeton Microarray Facility) and yielded ∼1 million 150 nucleotide paired-end reads. Reads were mapped to gDNA and cDNA reference sequences using bwa-mem (Li 2013) and a Variant Calling File (VCF) file was produced using the Naive Variant Caller as implemented in Galaxy (Version 0.0.2). Allele counts at focal sites (104, 111 and 122) for both cDNA and gDNA were generated using a custom script. ASE was estimated using the Cochran-Mantel-Haenzel framework as implemented in R (mantelhaen.test). Specifically, ASE is estimated as the relative risk of the mutant substitution in the cDNA population using allele counts from gDNA as a reference population.

### Behavioral phenotyping

We quantified “bang sensitivity” which is a measure of susceptibility to seizures and paralysis upon mechanical over-stimulation (Ganetzky and Wu 1982). As such, the bang sensitivity phenotype is a measure of proper neuron function. Individual flies were placed in an empty fly media vial and vortexed at the maximum setting for 20 seconds. Immediately following mechanical overstimulation, neurologically dysfunctional flies typically experience a period of convulsions and seizures. The recovery time was recorded as the time for each fly to right itself. Male flies were assayed 14 days post-eclosion. Approximate 95% confidence intervals were estimated around means by bootstrap resampling with replacement. An average of 43 flies were assayed (range 37-85). Recovery time distributions were compared using a Wilcoxon two-sample rank sum test with continuity correction as implemented in R (wilcox.test).

### CG exposure assay

As a measure of the ability to tolerate CGs, we exposed adult flies (1-7 days post-eclosion) to media containing known concentrations of the CG ouabain (Sigma). 1.5 grams of Drosophila instant media (Carolina Biological Supply) was reconstituted in a plastic vial with 7 mL of either 0, 5, 10 or 20 mM ouabain. After food solidified (30 min), a small paper wick was added. Three replicates of 10 males and 10 females were kept in each vial at 25°C and 50% humidity. Mortality was measured after 7 days. The genetically engineered wild-type line was used as a control. The Cochran-Mantel-Haenzel test framework (implemented in R) was used to assess significant differences between treatments (i.e. 5/10/20 mM ouabain versus no ouabain), as well as estimates and 95% confidence bounds for the relative risk associated with treatment.

Odds Ratios (*OR*) estimated using this framework were converted into relative risk (*RR*) estimates using the formula, 55 = 65/(1 − 8 + (8 ∗ 65)), where *p* is the risk in the no ouabain control group. A constant (0.5) was added to all cells to allow for calculation of relative risk in cases where mortality or survivors were absent. Thus, the maximum relative risk for this sample size (three replicates of 20 individuals each) is limited to 117. We found no sex differences in survival.

## Acknowledgements

We thank M. Przeworski and M. Schumer for comments on the manuscript, D. James and A. Wilson for sending *Syntomeida epilais* specimens and M. Aguadé for providing pre-publication access to the *D. guanche* ATPα1 sequence. We thank Sabrina Stiehler and Miyoung Yang for technical assistance. Funding: This work was funded by NIH R01 GM115523 to PA, NIH T32 GM008424 to BPR, NIH R01 GM108073 and NIH R01 AG027453 to MJP, and DFG PE 2059/3-1 and the LOEWE program of the State of Hesse (Insect Biotechnology & Bioresources), Germany to GP. Author contributions: PA, AMT, LY, and MJP designed the study; All authors contributed to the experimental work. PA wrote the paper. AMT, LY, AP, BPR, ADT, GP, and MJP reviewed and edited the manuscript.

---

**Supplementary Figure S1.**
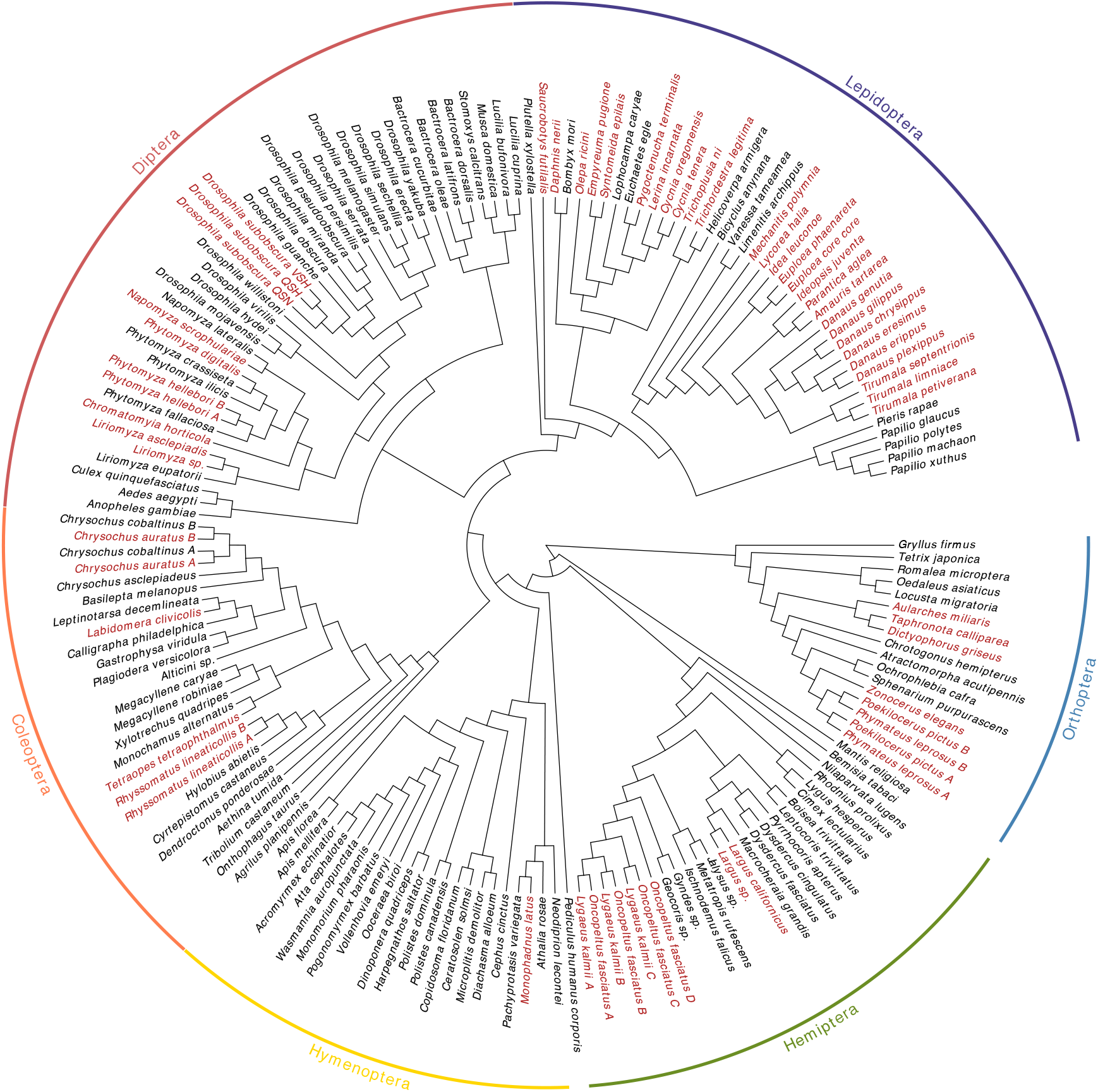
Phylogeny showing relationships of sampled species. In red are species known specialists on CG-containing plants or inferred to be CG-insensitive.

**Supplementary Figure S2.**
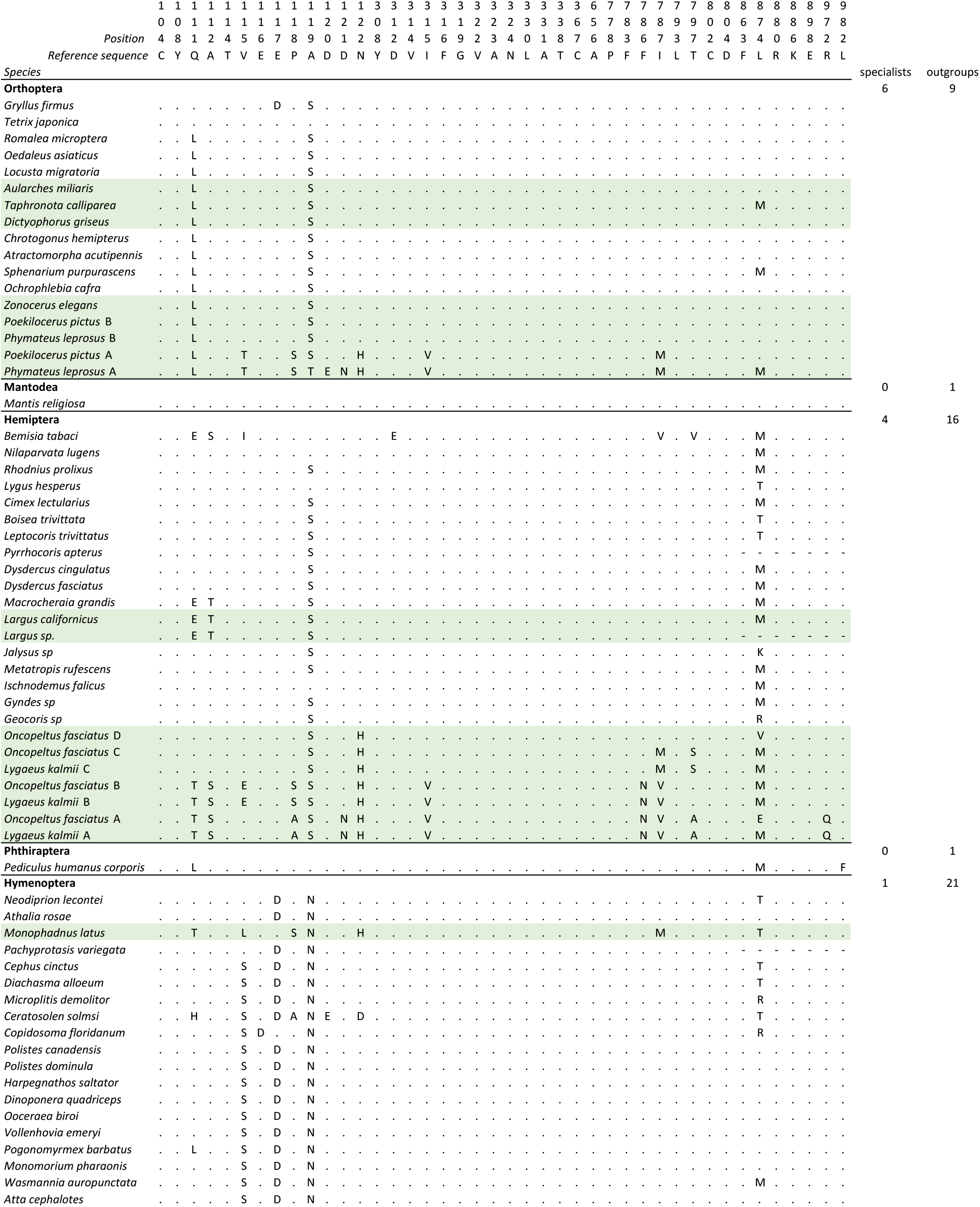
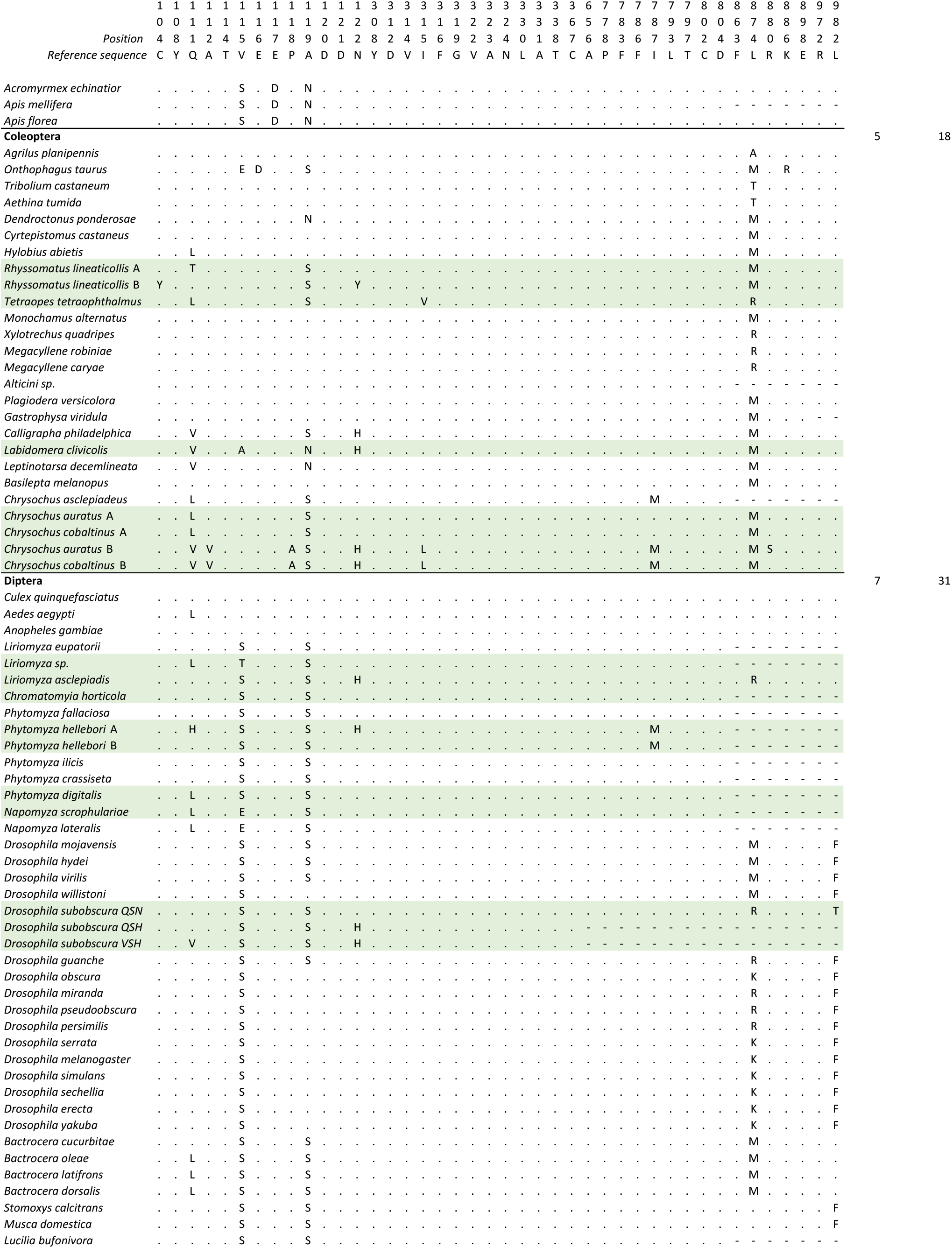
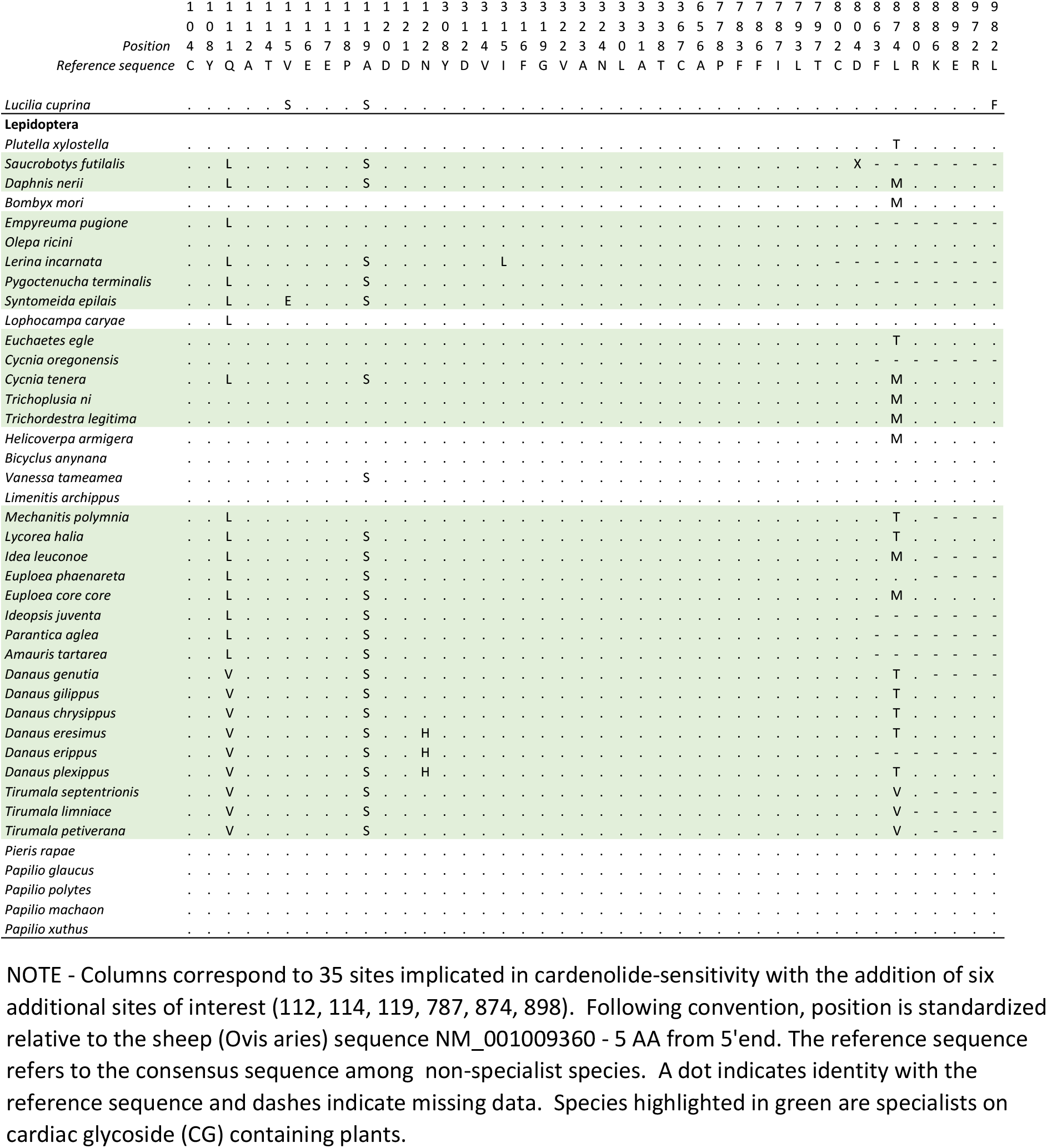
Summary of amino acid variation at sites implicated in cardenolide-sensitivity in ATP⍺1 in surveyed species

**Supplementary Figure S3.**
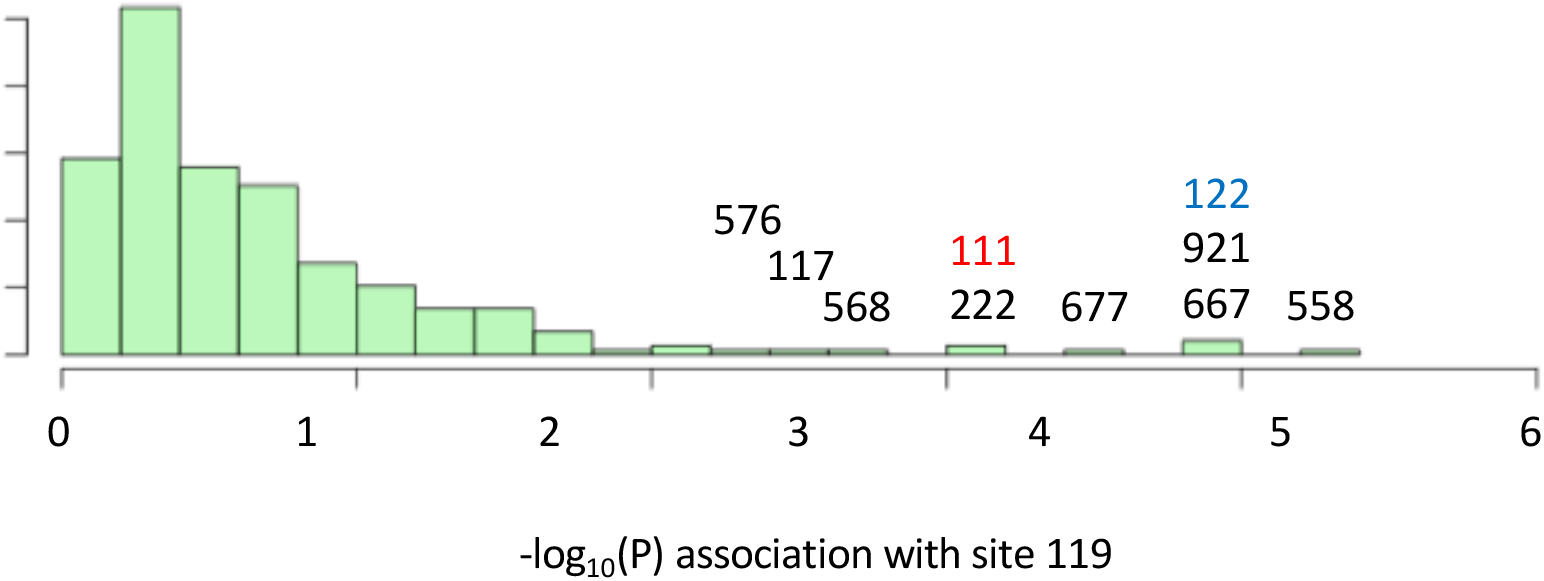
Variant sites in ATPα1 most strongly correlated with substitutions with site 119. Shown is the distribution of P-values for strength of phylogenetic correlation between site 119 and 270 non-singleton amino acid variants in an alignment including 174 ATPα1 sequences representing 161 species See Methods for a description of estimating correlations using BayesTraits. Indicated are the 10 top ranking sites. Sites 111 (red) and 122 (blue) are highly-correlated interacting sites known to be important for insensitivity of the Na^+^,K^+^-ATPase to CG-inhibition (see Figure 1).

**Supplementary Figure S4.**
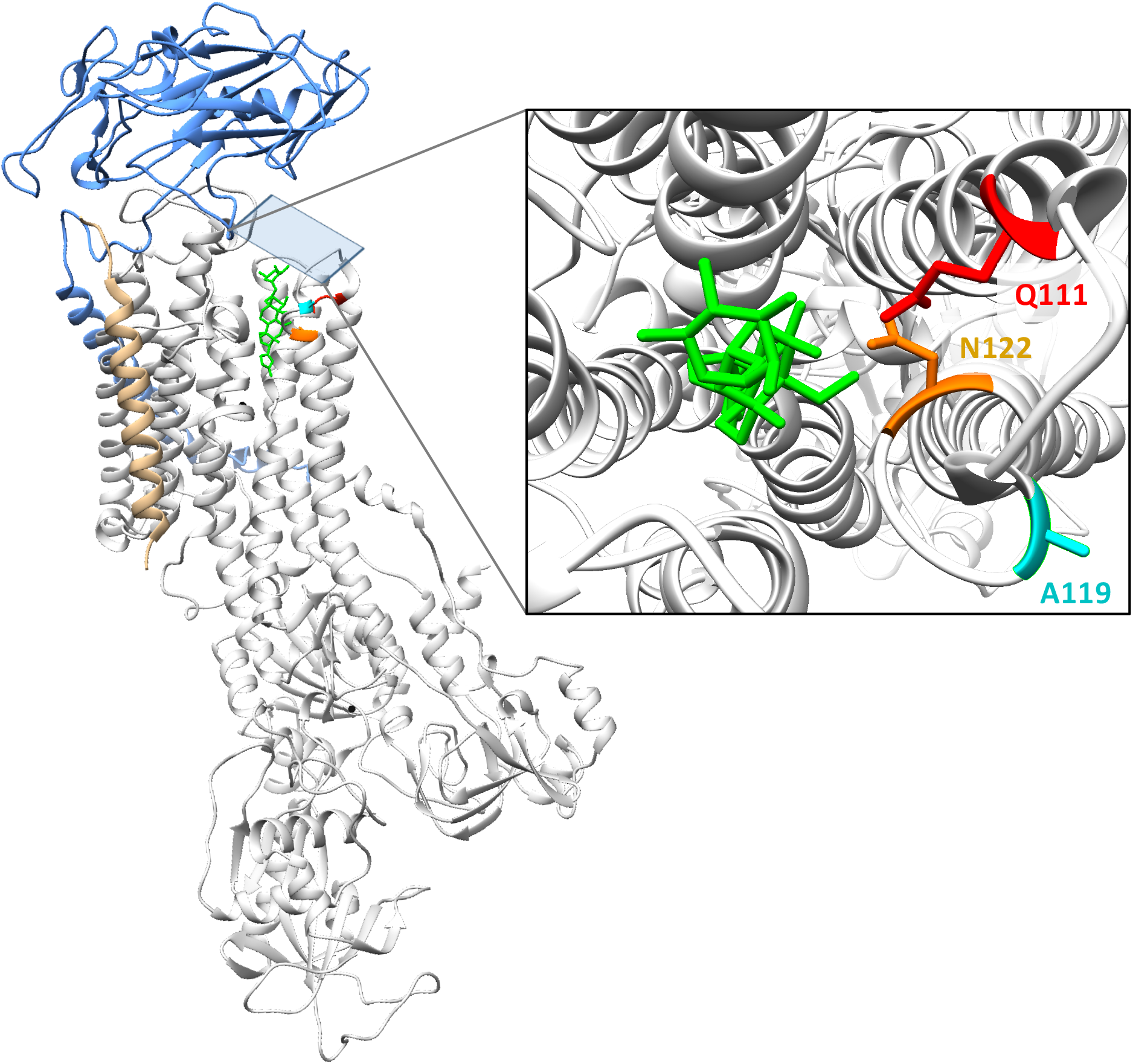
Estimated co-crystal structure of *Drosophila melanogaster* Na^+^,K^+^-ATPase bound to the cardiac glycoside (CG) ouabain. A homology model was constructed with Swiss-Model (Waterhouse et al. 2018; https://swissmodel.expasy.org), using the *Sus scrofa* (pig) structure bound to ouabain (PDB:4HYT) as a template and ATPα1 of *D. melanogaster* (Genbank: P13607) as target. The alpha-subunit is shown in white ribbon, and subunits beta and gamma in blue and gold ribbon, respectively. In red, magenta and orange are residues Q111, A119 and N122, respectively. The bound ouabain ligand is colored green. The structure was visualized using UCSF Chimera 1.11.2.

**Supplementary Figure S5.**
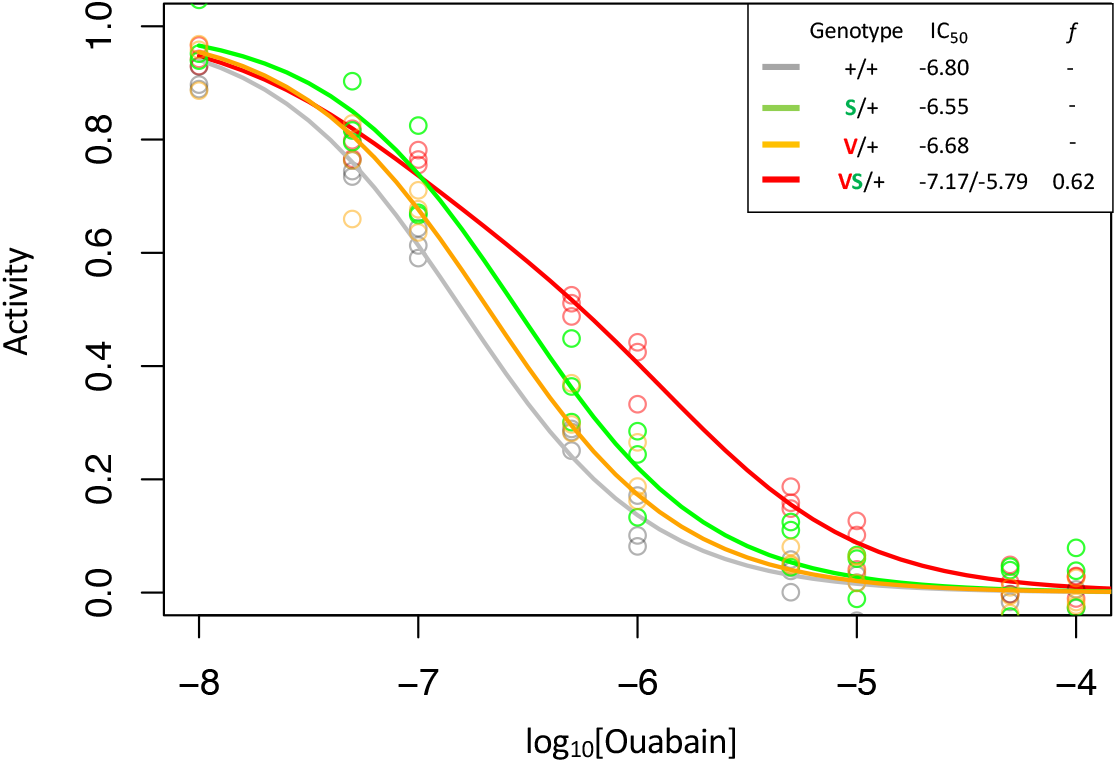
Enzyme inhibition curves for Q111V (V) with and without A119S (S). Following Figure 2, inhibition curves are plotted for heterozygous individuals to allow comparison of the effects of these substitutions in the presence and absence of A119S. Points represent biological replicates, which is the means across technical replicates. Curves for the engineered wild-type strain (+/+) and A119S/+ are plotted for comparison. The presence of A119S or Q111V individually exhibit monophasic curves suggesting small effects on CG-insensitivity. In contrast, Q111V in combination with A119S (VS/+) results in a right-shifted curve indicating increased CG-insensitivity (IC_50,2_ = 1.6e-6). For VS/+, the 95% CI on the estimated proportion of CG-inhibitable activity of the mutant form, *f* = 0.54-0.69.

**Supplementary Figure S6.**
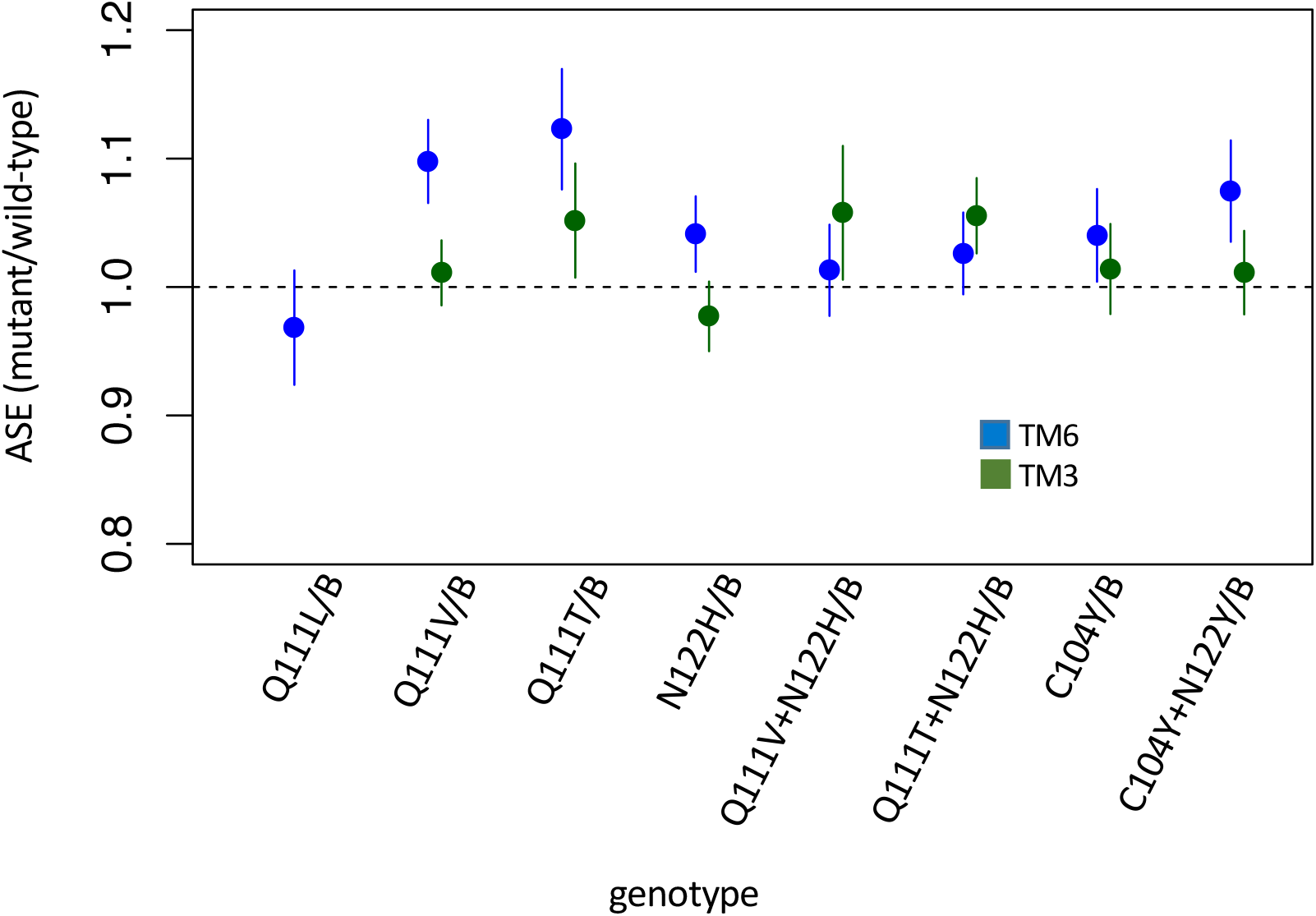
Allele-specific expression (ASE) in heterozygous lines. Estimates and 95% confidence intervals are based on Cochran-Mantel-Haenzel framework analysis – i.e. plotted is “relative risk” of the mutant allele in the cDNA population (treatment) using genomic DNA as a control population (see Methods). Equal expression of the two alleles corresponds to a value of 1. Estimates and confidence intervals are shown for alleles balanced over two standard balancer chromosomes, B = TM6 (blue) or TM3 (green) which carries a wild-type allele of ATPα1. While significant deviations from equal expression (correcting for multiple comparisons) are observed for Q111V/TM6, Q111T/TM6, C104Y+N122Y/TM6, Q111T+N122H/TM3, the magnitude of the differences are small and in favor of the mutant allele.

**Supplementary Figure S7.**
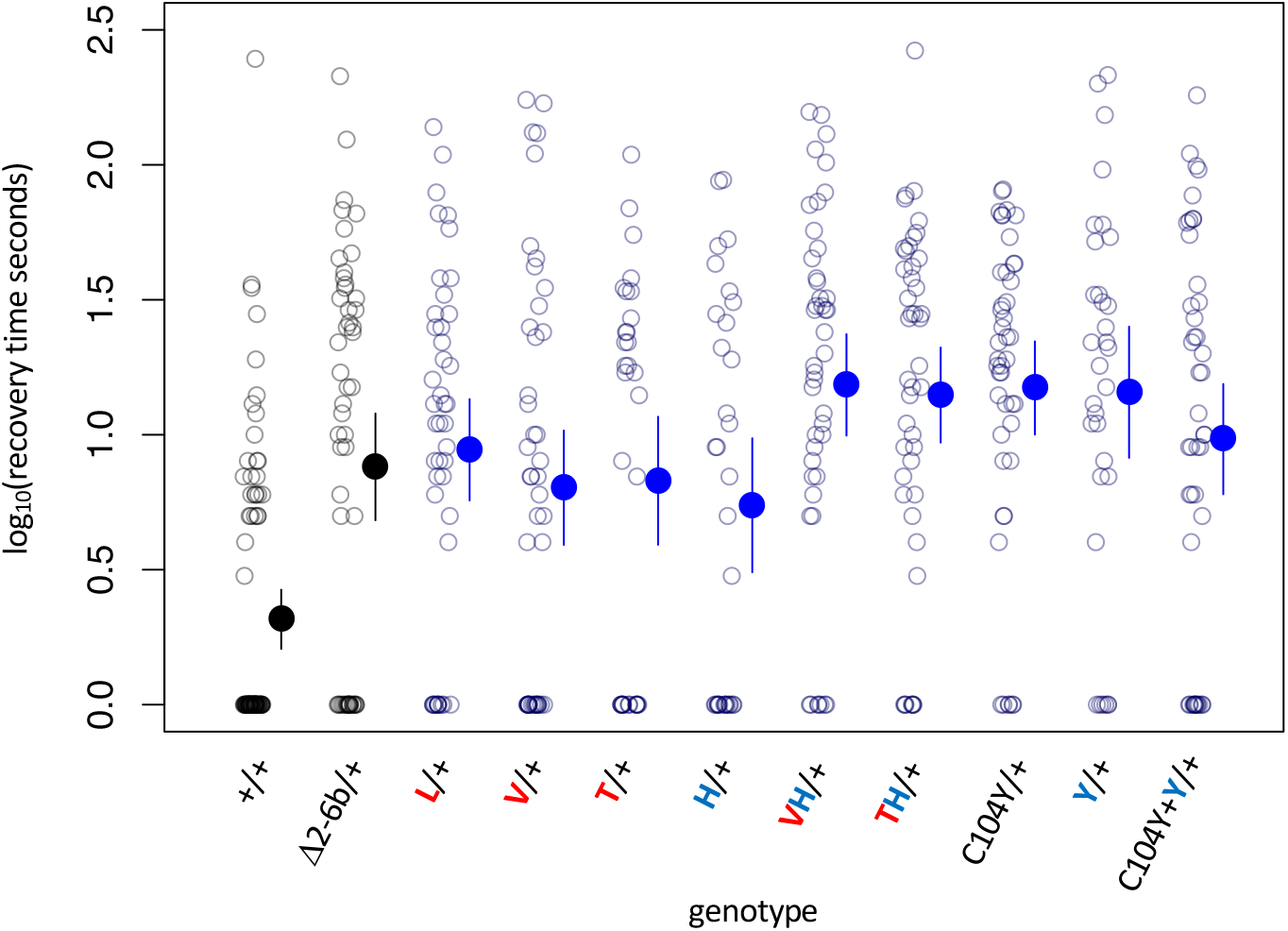
Bang sensitivity phenotypes of lines with heterozygous substitutions at sites 104, 111 and 122. Δ2-6b is a loss of function deletion of ATPα1. Red letters indicate substitutions at position 111; blue letters indicate substitutions at position 122. Labels: + = wild-type; Δ2-6b = loss of function deletion of exons 2 through 6b; L = Q111L; V = Q111V; T = Q111T; H = N122H; Y=N122Y. All substitutions are heterozygous over the engineered wild-type allele (+). In all cases, the distributions for substitution lines (Mut/+) are significantly different than the engineered wild-type strain (+/+) (*P* < 5e-5, Wilcoxon test with continuity correction, correcting for multiple tests).

**Supplementary Figure S8.**
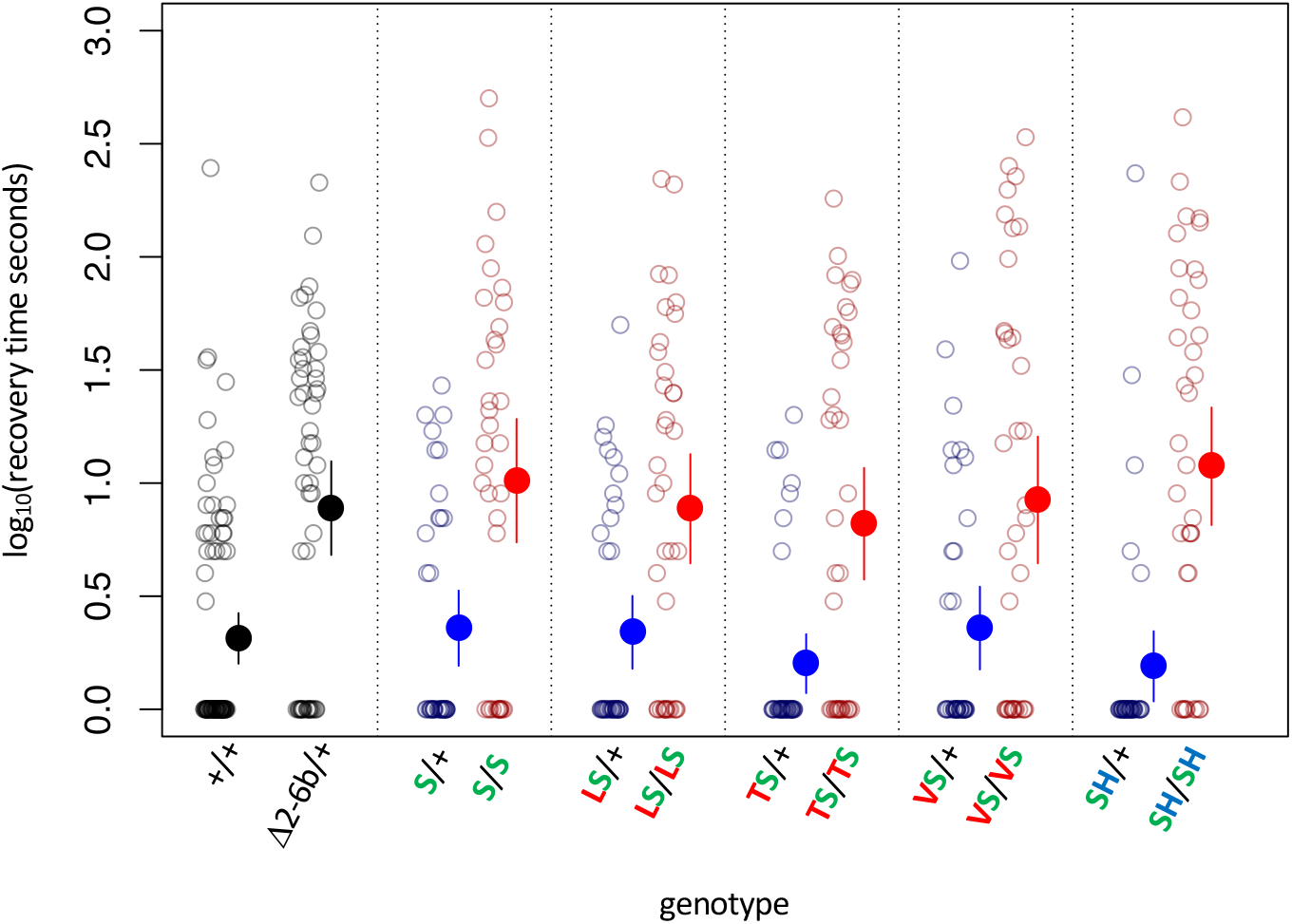
Bang sensitivity phenotypes of substitutions at sites 111 and 122 on the background of A119S. Δ2-6b is a loss of function deletion of ATPα1. Red letters indicate substitutions at position 111; green letters at position 119; blue letters at position 122. Blue points and whiskers indicate substitutions that are heterozygous for wild-type allele (+). Red points and whiskers indicate data for homozygous substitutions. Labels: + = engineered wild-type allele; Δ2-6 = loss of function deletion of exons 2-6b; S = A119S; LS = Q111L+A119S; VS = Q111V+A119S; TS = Q111T+A119S; SH = A119S+N122H. In all cases, the distributions for heterozygotes (Mut/+) and homozygotes (Mut/Mut) are significantly different (*P*<0.005, Wilcoxon test with continuity correction, corrected for multiple tests).

**Supplementary Figure S9.**
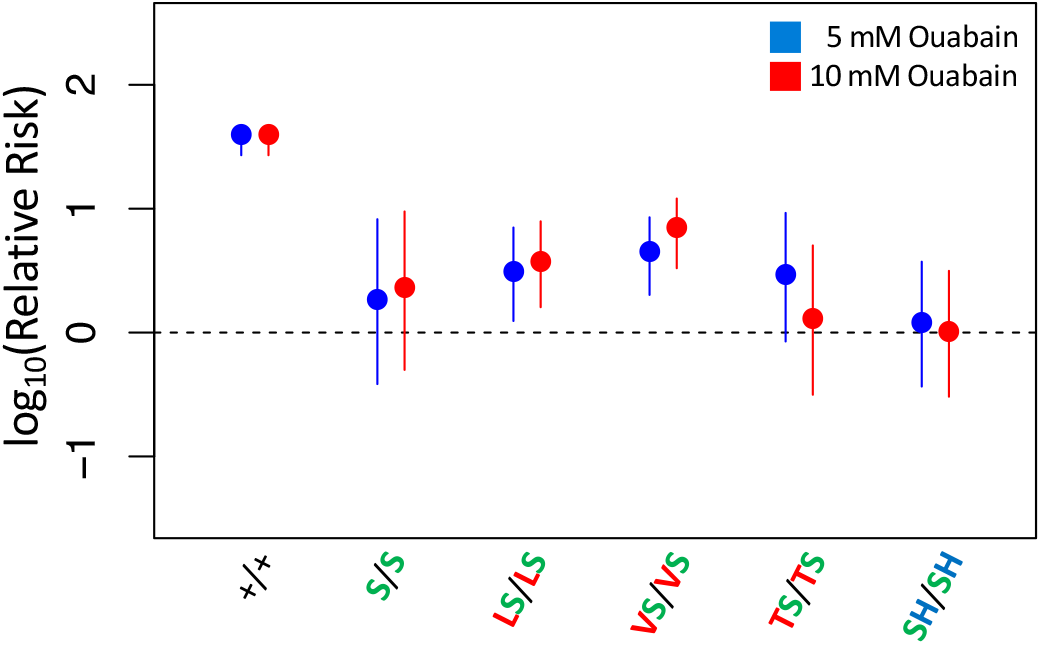
Adult survival of homozygous strains with A119S upon 7-day exposure to CGs. Plotted is the log relative risk of treatment, for 5 mM (blue) or 10 mM (red) ouabain, relative to no treatment controls (no ouabain) for homozygous strains. Points and whiskers represent estimates and 95% confidence bounds estimated using the Cochran-Mantel-Haenzel framework (Methods). Each estimate is based in three biological replicates of 20 flies per concentration. A value of 0 corresponds to equal probability of survival of treatment versus control indicating complete insensitivity to the tested ouabain concentration. Labels: +/+ = engineered wild-type strain; S/S = A119S homozygote; LS/LS = Q111L+A119S homozygote; VS/VS = Q111V+A119S homozygote; TS/TS = Q111T+A119S homozygote; SH/SH = A119S+N122H homozygote.

**Supplementary Figure S10.**
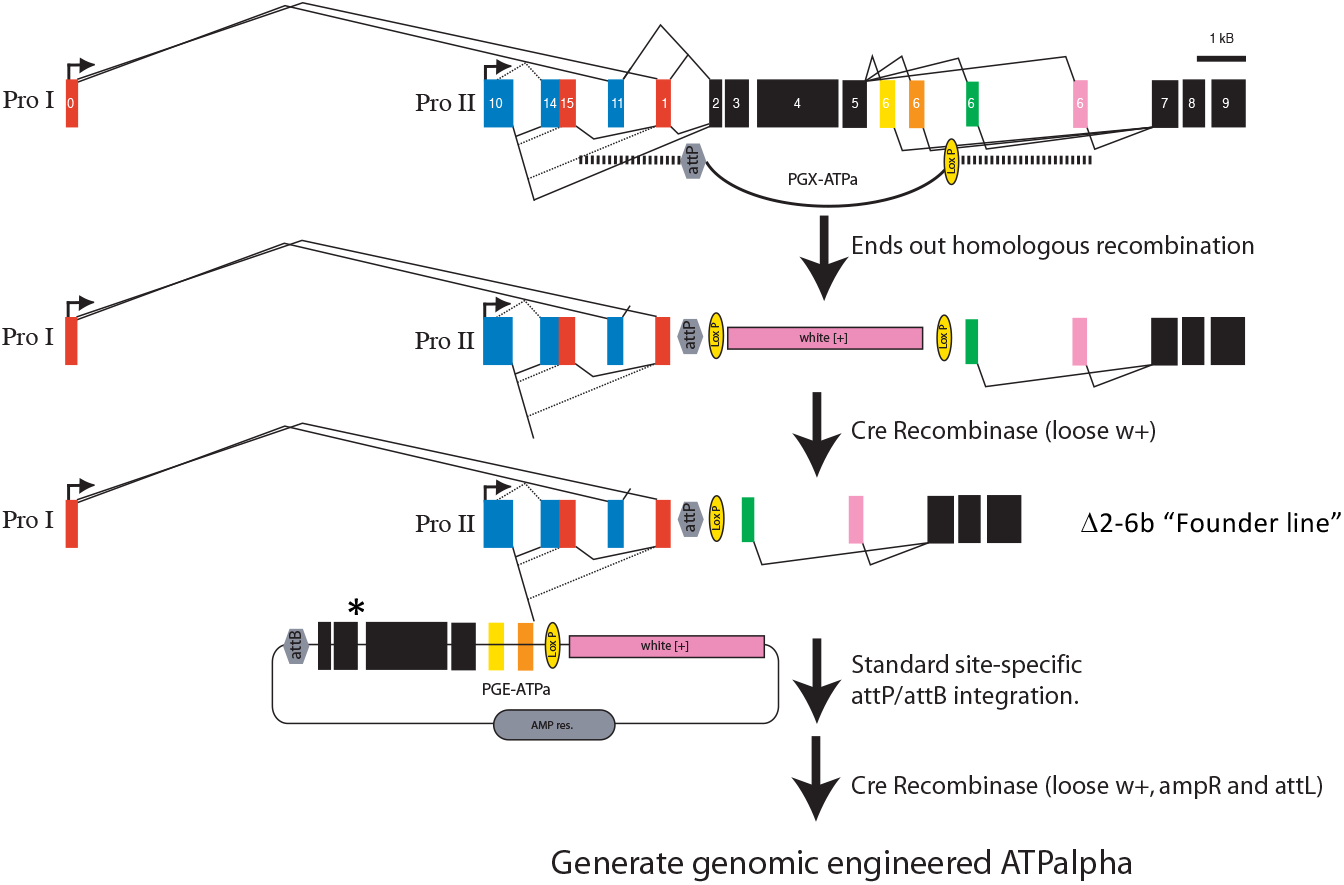
Engineering strategy for generating amino acid substitution lines. A targeted replacement of exons 2-6b with a mini-white gene was generated by ends-out homologous recombination. The mini-white gene was removed using cre-lox site-specific recombination to create the Δ2-6b “Founder line”. The founder line is injected with the PGE-ATPa plasmid carrying a substitution of choice (asterisk) and incorporated by site-specific phiC31/attP integration. The mini-white gene was removed using cre-lox site-specific recombination to create engineered ATPα (ATPalpha) lines.

**Supplementary Table S1.** Sources of sequence data used in this study.

| Species | Order | Method | Accession | Reference or BioProject |
| --- | --- | --- | --- | --- |
| *Gryllus firmus* | Orthoptera | *RNAseq de novo* assembly | Pending | Yang *et al.* 2019 |
| *Tetrix japonica* | Orthoptera | *RNAseq de novo* assembly | Pending | Yang *et al.* 2019 |
| *Romalea microptera* | Orthoptera | *RNAseq de novo* assembly | Pending | Yang *et al.* 2019 |
| *Oedaleus asiaticus* | Orthoptera | *RNAseq de novo* assembly | Pending | Yang *et al.* 2019 |
| *Locusta migratoria* | Orthoptera | Sanger cDNA sequencing | KF813097.1 | Bao, 2013 |
| *Aularches miliaris* | Orthoptera | *RNAseq de novo* assembly | MK294065 | Yang *et al.* 2019 |
| *Taphronota calliparea* | Orthoptera | *RNAseq de novo* assembly | MK294066 | Yang *et al.* 2019 |
| *Dictyophorus griseus* | Orthoptera | *RNAseq de novo* assembly | MK294067 | Yang *et al.* 2019 |
| *Chrotogonus hemipterus* | Orthoptera | *RNAseq de novo* assembly | MK294068 | Yang *et al.* 2019 |
| *Atractomorpha acutipennis* | Orthoptera | *RNAseq de novo* assembly | MK294069 | Yang *et al.* 2019 |
| *Ochrophlebia cafra* | Orthoptera | *RNAseq de novo* assembly | MK294070 | Yang *et al.* 2019 |
| *Sphenarium purpurascens* | Orthoptera | *RNAseq de novo* assembly | MK294071 | Yang *et al.* 2019 |
| *Zonocerus elegans* | Orthoptera | *RNAseq de novo* assembly | MK294072 | Yang *et al.* 2019 |
| *Poecilocerus pictus B* | Orthoptera | *RNAseq de novo* assembly | MK294074 | Yang *et al.* 2019 |
| *Phymateus leprosus B* | Orthoptera | *RNAseq de novo* assembly | MK294073 | Yang *et al.* 2019 |
| *Poecilocerus pictus A* | Orthoptera | *RNAseq de novo* assembly | MK294076 | Yang *et al.* 2019 |
| *Phymateus leprosus A* | Orthoptera | *RNAseq de novo* assembly | MK294075 | Yang *et al.* 2019 |
| *Mantis religiosa* | Mantodea | *RNAseq de novo* assembly | Pending | Yang *et al.* 2019 |
| *Bemisia tabaci* | Hemiptera | Query reference genome | XM_019041063.1 | PRJNA352527 |
| *Nilaparvata lugens* | Hemiptera | Query reference genome | XM_022328639.1 | PRJNA398259 |
| *Rhodnius prolixus* | Hemiptera | Query reference genome | KQ034075.1 | PRJNA13648 |
| *Lygus hesperus* | Hemiptera | *RNAseq de novo* assembly | GBRD01003681.1 | PRJNA210219 |
| *Cimex lectularius* | Hemiptera | Query reference genome | XM_014394884.2 | PRJNA298750 |
| *Boisea trivittata* | Hemiptera | *RNAseq de novo* assembly | JQ771499.1 | Zhen *et al.* 2012 |
| *Leptocoris trivittatus* | Hemiptera | *RNAseq de novo* assembly | JQ771499.1 | Zhen *et al.* 2012 |
| *Pyrrhocoris apterus* | Hemiptera | PCR-sequencing cDNA | HE956739.1 | Dobler *et al.* 2012 |
| *Dysdercus cingulatus* | Hemiptera | *RNAseq de novo* assembly | Pending | SRR5040256 |
| *Dysdercus fasciatus* | Hemiptera | *RNAseq de novo* assembly | Pending | SRR489295 |
| *Macrocheraia grandis* | Hemiptera | *RNAseq de novo* assembly | Pending | SRR5040251 |
| *Largus californicus* | Hemiptera | *RNAseq de novo* assembly | Pending | SRR1821933 |
| *Largus sp.* | Hemiptera | PCR-sequencing cDNA | HE956738.1 | Dobler *et al.* 2012 |
| *Jalysus sp.* | Hemiptera | *RNAseq de novo* assembly | Pending | SRR1821926 |
| *Metatropis rufescens* | Hemiptera | *RNAseq de novo* assembly | Pending | SRR2051503 |
| *Ischnodemus falicus* | Hemiptera | *RNAseq de novo* assembly | Pending | SRR1821925 |
| *Gyndes sp.* | Hemiptera | *RNAseq de novo* assembly | Pending | SRR5137185 |
| *Geocoris sp.* | Hemiptera | *RNAseq de novo* assembly | Pending | SRR1821921 |
| *Oncopeltus fasciatus D* | Hemiptera | *RNAseq de novo* assembly | MK765670 | Yang *et al.* 2019 |
| *Oncopeltus fasciatus C* | Hemiptera | *RNAseq de novo* assembly | JQ771518.1 | Zhen *et al.* 2012 |
| *Lygaeus kalmii C* | Hemiptera | *RNAseq de novo* assembly | JQ771515.1 | Zhen *et al.* 2012 |
| *Oncopeltus fasciatus B* | Hemiptera | *RNAseq de novo* assembly | JQ771519.1 | Zhen *et al.* 2012 |
| *Lygaeus kalmii B* | Hemiptera | *RNAseq de novo* assembly | JQ771514.1 | Zhen *et al.* 2012 |
| *Oncopeltus fasciatus A* | Hemiptera | *RNAseq de novo* assembly | JQ771520.1 | Zhen *et al.* 2012 |
| *Lygaeus kalmii A* | Hemiptera | *RNAseq de novo* assembly | JQ771513.1 | Zhen *et al.* 2012 |
| *Pediculus humanus corporis* | Phthiraptera | Query reference genome | XM_002427669.1 | Kirkness *et al.* 2007 |
| *Neodiprion lecontei* | Hymenoptera | Query reference genome | XM_015658649.1 | PRJNA312506 |
| *Athalia rosae* | Hymenoptera | Query reference genome | XM_012414228.2 | PRJNA282653 |
| *Monophadnus latus* | Hymenoptera | *RNAseq de novo* assembly | Pending | This study |
| *Pachyprotasis variegata* | Hymenoptera | PCR-sequencing cDNA | LN736263.1 | Dobler *et al.* 2015 |
| *Cephus cinctus* | Hymenoptera | Query reference genome | XM_015729628.2 | PRJNA297591 |
| *Diachasma alloenum* | Hymenoptera | Query reference genome | XM_015265214.1 | PRJNA306876 |
| *Microplitis demolitor* | Hymenoptera | Query reference genome | XM_008550885.1 | PRJNA251518 |
| *Ceratosolen solmsi* | Hymenoptera | Query reference genome | XM_011505654.1 | PRJNA277475 |
| *Copidosoma floridanum* | Hymenoptera | Query reference genome | XM_014350407.2 | PRJNA297581 |
| *Polistes canadensis* | Hymenoptera | Query reference genome | XM_014749330.1 | PRJNA301748 |
| *Polistes dominula* | Hymenoptera | Query reference genome | XM_015323700.1 | PRJNA307991 |
| *Harpegnathos saltator* | Hymenoptera | Query reference genome | XM_025304334.1 | PRJNA476946 |
| *Dinoponera quadricaps* | Hymenoptera | Query reference genome | XM_014620740.1 | PRJNA301625 |
| *Ooceraea biroi* | Hymenoptera | Query reference genome | XM_011345351.3 | PRJNA501908 |
| *Vollenhovia emeryi* | Hymenoptera | Query reference genome | XM_012020948.1 | PRJNA278668 |
| *Pogonomyrmex barbatus* | Hymenoptera | Query reference genome | XM_011645049.2 | PRJNA276107 |
| *Monomorium pharaonis* | Hymenoptera | Query reference genome | XM_012678519.2 | PRJNA479782 |
| *Wasmannia auropunctata* | Hymenoptera | Query reference genome | XM_011708535.1 | PRJNA279179 |
| *Atta cephalotes* | Hymenoptera | Query reference genome | XM_012202049.1 | PRJNA279976 |
| *Acromyrmex echinator* | Hymenoptera | Query reference genome | XM_011052414.1 | PRJNA271903 |
| *Apis mellifera* | Hymenoptera | Query reference genome | XM_020018205.1 | PRJNA361278 |
| *Apis florea* | Hymenoptera | Query reference genome | XM_012490248.1 | PRJNA86991 |
| *Agrilus planipennis* | Coleoptera | Query reference genome | XM_018472182.2 | PRJNA343475 |
| *Onthophagus taurus* | Coleoptera | Query reference genome | XM_023047807.1 | PRJNA419349 |
| *Tribolium castaneum* | Coleoptera | Query reference genome | XM_008198195.2 | PRJNA15718 |
| *Aethina tumida* | Coleoptera | Query reference genome | XM_020018205.1 | PRJNA361278 |
| *Dendroctonus ponderosae* | Coleoptera | Query reference genome | XM_019913697.1 | PRJNA360270 |
| *Cyrtepidomus castaneus* | Coleoptera | *RNAseq de novo* assembly | JQ771502.1 | Zhen *et al.* 2012 |
| *Hylobius abietis* | Coleoptera | *RNAseq de novo* assembly | Pending | SRR6765939 |
| *Rhyssomatus lineaticollis A* | Coleoptera | *RNAseq de novo* assembly | JQ771524.1 | Zhen *et al.* 2012 |
| *Rhyssomatus lineaticollis B* | Coleoptera | *RNAseq de novo* assembly | JQ771523.1 | Zhen *et al.* 2012 |
| *Tetraopes tetraophthalmus* | Coleoptera | *RNAseq de novo* assembly | JQ771526.1 | Zhen *et al.* 2012 |
| *Monochamus alternatus* | Coleoptera | *RNAseq de novo* assembly | Pending | SRR2521326 |
| *Xylotrechus quadripes* | Coleoptera | *RNAseq de novo* assembly | Pending | SRR7077078 |
| *Megacyllene robiniae* | Coleoptera | *RNAseq de novo* assembly | JQ771517.1 | Zhen *et al.* 2012 |
| *Megacyllene caryae* | Coleoptera | *RNAseq de novo* assembly | Pending | SRR1586007 |
| *Alticini sp.* | Coleoptera | PCR-sequencing cDNA | HE956742.1 | Dobler *et al.* 2012 |
| *Plagiodera versicolora* | Coleoptera | *RNAseq de novo* assembly | JQ771522.1 | Zhen *et al.* 2012 |
| *Gastrophysa viridula* | Coleoptera | PCR-sequencing cDNA | HE956744.1 | Dobler *et al.* 2012 |
| *Calligrapha philadelphia* | Coleoptera | *RNAseq de novo* assembly | Pending | ERR1333730 |
| *Labidomera clivicollis* | Coleoptera | *RNAseq de novo* assembly | JQ771511.1 | Zhen *et al.* 2012 |
| *Leptinotarsa decemlineata* | Coleoptera | Query reference genome | XM_023161024.1 | PRJNA420356 |
| *Basilepta melanopus* | Coleoptera | *RNAseq de novo* assembly | Pending | SRR8187395 |
| *Chrysochus asclepiadeus* | Coleoptera | PCR-sequencing cDNA | HE956740.1 | Dobler *et al.* 2012 |
| *Chrysochus auratus A* | Coleoptera | *RNAseq de novo* assembly | JQ771500.1 | Zhen *et al.* 2012 |
| *Chrysochus cobalitinus A* | Coleoptera | *RNAseq de novo* assembly | MK765671 | Yang *et al.* 2019 |
| *Chrysochus auratus B* | Coleoptera | *RNAseq de novo* assembly | JQ771501.1 | Zhen *et al.* 2012 |
| *Chrysochus cobalitinus B* | Coleoptera | *RNAseq de novo* assembly | MK765672 | Yang *et al.* 2019 |
| *Culex quinquefasciatus* | Diptera | Query reference genome | NW_001886804.1 | PRJNA29017 |
| *Aedes aegypti* | Diptera | Query reference genome | NC_035107.1 | PRJNA318737 |
| *Anopheles gambiae* | Diptera | Query reference genome | XM_020076988.1 | PRJNA357111 |
| *Liriomyza eupatorii* | Diptera | PCR-sequencing cDNA | LT795109.1 | Petschenka *et al.* 2017 |
| *Liriomyza sp.* | Diptera | PCR-sequencing cDNA | HE956748.1 | Dobler *et al.* 2012 |
| *Liriomyza asclepiadis* | Diptera | *RNAseq de novo* assembly | MK294077 | Yang *et al.* 2019 |
| *Chromatomyia horticola* | Diptera | PCR-sequencing cDNA | LT795081.1 | Petschenka *et al.* 2017 |
| *Phytomyza fallaciosa* | Diptera | PCR-sequencing cDNA | LT795082.1 | Petschenka *et al.* 2017 |
| *Phytomyza hellebori A* | Diptera | PCR-sequencing cDNA | LT795110.1 | Petschenka *et al.* 2017 |
| *Phytomyza hellebori B* | Diptera | PCR-sequencing cDNA | LT795111.1 | Petschenka *et al.* 2017 |
| *Phytomyza crassiseti* | Diptera | PCR-sequencing cDNA | LT795078.1 | Petschenka *et al.* 2017 |
| *Phytomyza ilicis* | Diptera | PCR-sequencing cDNA | LT795077.1 | Petschenka *et al.* 2017 |
| *Phytomyza digitalis* | Diptera | PCR-sequencing cDNA | LT795083.1 | Petschenka *et al.* 2017 |
| *Napomyza scrophulariae* | Diptera | PCR-sequencing cDNA | LT795080.1 | Petschenka *et al.* 2017 |
| *Napomyza lateralis* | Diptera | PCR-sequencing cDNA | LT795079.1 | Petschenka *et al.* 2017 |
| *Drosophila mojavensis* | Diptera | Query reference genome | XM_015167953.1 | PRJNA29977 |
| *Drosophila hydei* | Diptera | Query reference genome | XM_023321093.1 | PRJNA422293 |
| *Drosophila virilis* | Diptera | Query reference genome | XM_002055964.2 | PRJNA29995 |
| *Drosophila willistoni* | Diptera | Query reference genome | XM_015177397.2 | PRJNA29997 |
| *Drosophila subobscura* QSN | Diptera | *RNAseq de novo* assembly | Pending | This study |
| *Drosophila subobscura* QSH | Diptera | PCR-sequencing gDNA | KT318950.1 | Pegueroles *et al.* 2016 |
| *Drosophila subobscura* VSH | Diptera | PCR-sequencing cDNA | KT318946.1 | Pegueroles *et al.* 2016 |
| *Drosophila guanche* | Diptera | Query reference genome | Pending | Puerma *et al.* 2018 |
| *Drosophila obscura* | Diptera | Query reference genome | XM_022353228.1 | PRJNA399719 |
| *Drosophila miranda* | Diptera | Query reference genome | XM_017286564.1 | PRJNA325520 |
| *Drosophila pseudoobscura* | Diptera | Query reference genome | XM_001358575.3 | Drosophila 12 Genomes Consortium, 2007 |
| *Drosophila persimilis* | Diptera | Query reference genome | XM_026985943.1 | PRJNA501994 |
| *Drosophila serrata* | Diptera | Query reference genome | XM_020943893.1 | PRJNA384652 |
| *Drosophila melanogaster* | Diptera | Query reference genome | FBgn0002921 | ftp://ftp.flybase.net/releases/FB2019_01/ |
| *Drosophila simulans* | Diptera | Query reference genome | XM_002102615.2 | PRJNA297806 |
| *Drosophila sechellia* | Diptera | Query reference genome | XM_002044256.1 | Drosophila 12 Genomes Consortium, 2007 |
| *Drosophila erecta* | Diptera | Query reference genome | XM_001979324.3 | PRJNA501994 |
| *Drosophila yakuba* | Diptera | Query reference genome | XM_015191921.1 | PRJNA29999 |
| *Bactrocera cucurbitae* | Diptera | Query reference genome | XM_011190816.1 | PRJNA273817 |
| *Bactrocera oleae* | Diptera | Query reference genome | XM_014232491.1 | PRJNA293367 |
| *Bactrocera latifrons* | Diptera | Query reference genome | XM_018929031.1 | PRJNA351211 |
| *Bactrocera dorsalis* | Diptera | Query reference genome | XM_011214825.2 | PRJNA273958 |
| *Stomoxys calcitrans* | Diptera | Query reference genome | XM_013260537.1 | PRJNA288986 |
| *Musca domestica* | Diptera | Query reference genome | XM_020037539.1 | PRJNA210139 |
| *Lucilia bufonivora* | Diptera | PCR-sequencing cDNA | HG938131.1 | Mebs *et al.* 2014 |
| *Lucilia cuprina* | Diptera | Query reference genome | XM_023439570.1 | PRJNA423280 |
| *Plutella xylostella* | Lepidoptera | Query reference genome | XM_011549930.1 | PRJNA277936 |
| *Saucrobotys futilalis* | Lepidoptera | PCR-sequencing cDNA | HE956749.1 | Dobler *et al.* 2012 |
| *Daphnis nerii* | Lepidoptera | *RNAseq de novo* assembly | MK294081 | Yang *et al.* 2019 |
| *Bombyx mori* | Lepidoptera | cDNA sequencing | LC029030.1 | Homareda and Hara, unpublished |
| *Empyreuma pugione* | Lepidoptera | PCR-sequencing cDNA | LN736266.1 | Dobler *et al.* 2015 |
| *Olepa ricini* | Lepidoptera | *RNAseq de novo* assembly | MK294080 | Yang *et al.* 2019 |
| *Lerina incarnata* | Lepidoptera | PCR-sequencing cDNA | HE956754.1 | Dobler *et al.* 2012 |
| *Pygoctenucha terminalis* | Lepidoptera | PCR-sequencing cDNA | HE956753.1 | Dobler *et al.* 2012 |
| *Syntomeida epilais* | Lepidoptera | *RNAseq de novo* assembly | Pending | This study |
| *Lophocampa caryae* | Lepidoptera | *RNAseq de novo* assembly | JQ771510.1 | Zhen *et al.* 2012 |
| *Euchaetes egle* | Lepidoptera | *RNAseq de novo* assembly | JQ771508.1 | Zhen *et al.* 2012 |
| *Cynia oregonensis* | Lepidoptera | PCR-sequencing cDNA | HE956750.1 | Dobler *et al.* 2012 |
| *Cynia tenera* | Lepidoptera | *RNAseq de novo* assembly | JQ771504.1 | Zhen *et al.* 2012 |
| *Trichoplusia ni* | Lepidoptera | Query reference genome | XM_026879049.1 | PRJNA497582 |
| *Trichordestra legitima* | Lepidoptera | *RNAseq de novo* assembly | JQ771525.1 | Zhen *et al.* 2012 |
| *Helicoverpa armigera* | Lepidoptera | Query reference genome | XM_021340407.1 | PRJNA388211 |
| *Bicyclus anynana* | Lepidoptera | Query reference genome | XM_024099163.1 | PRJNA434100 |
| *Vanessa tameamea* | Lepidoptera | Query reference genome | XM_026633901.1 | PRJNA493654 |
| *Limenitis archippus* | Lepidoptera | *RNAseq de novo* assembly | JQ771509.1 | Zhen *et al.* 2012 |
| *Mechanitis polymnia* | Lepidoptera | PCR-sequencing cDNA | HF945460.1 | Petschenka *et al.* 2013 |
| *Lycorea halia* | Lepidoptera | *RNAseq de novo* assembly | JQ771512.1 | Zhen *et al.* 2012 |
| *Idea leuconoe* | Lepidoptera | PCR-sequencing cDNA | HF945457.1 | Petschenka *et al.* 2013 |
| *Euploea phaenareta* | Lepidoptera | PCR-sequencing cDNA | HF945456.1 | Petschenka *et al.* 2013 |
| *Euploea core core* | Lepidoptera | *RNAseq de novo* assembly | MK294079 | Yang *et al.* 2019 |
| *Ideopsis juvena* | Lepidoptera | PCR-sequencing cDNA | HF945458.1 | Petschenka *et al.* 2013 |
| *Parantica aglea* | Lepidoptera | PCR-sequencing cDNA | HF945461.1 | Petschenka *et al.* 2013 |
| *Amauris tartarea* | Lepidoptera | PCR-sequencing cDNA | HF945450.1 | Petschenka *et al.* 2013 |
| *Danaus genutia* | Lepidoptera | PCR-sequencing cDNA | HF945453.1 | Petschenka *et al.* 2013 |
| *Danaus gilippus* | Lepidoptera | *RNAseq de novo* assembly | JQ771506.1 | Zhen *et al.* 2012 |
| *Danaus chrysippus* | Lepidoptera | *RNAseq de novo* assembly | MK294078 | Yang *et al.* 2019 |
| *Danaus eresimus* | Lepidoptera | *RNAseq de novo* assembly | JQ771505.1 | Zhen *et al.* 2012 |
| *Danaus erippus* | Lepidoptera | PCR-sequencing cDNA | HF945451.1 | Petschenka *et al.* 2013 |
| *Danaus plexippus* | Lepidoptera | *RNAseq de novo* assembly | JQ771507.1 | Zhen *et al.* 2012 |
| *Tirumala septentrionis* | Lepidoptera | PCR-sequencing cDNA | HF945464.1 | Petschenka *et al.* 2013 |
| *Tirumala limniace* | Lepidoptera | PCR-sequencing cDNA | HF945462.1 | Petschenka *et al.* 2013 |
| *Tirumala petiverana* | Lepidoptera | PCR-sequencing cDNA | HF945463.1 | Petschenka *et al.* 2013 |
| *Pieris rapae* | Lepidoptera | Query reference genome | XM_022259192.1 | PRJNA397594 |
| *Papilio glaucus* | Lepidoptera | *RNAseq de novo* assembly | JQ771498.1 | Zhen *et al.* 2012 |
| *Papilio polytes* | Lepidoptera | Query reference genome | XM_013283743.1 | PRJNA291535 |
| *Papilio machaon* | Lepidoptera | Query reference genome | XM_014516575.1 | PRJNA300299 |
| *Papilio xuthus* | Lepidoptera | Query reference genome | XM_013315255.1 | PRJNA291600 |

**Supplementary Table S2.** References for the phylogenetic relationships of taxa used in this study.

| Group | References |
| --- | --- |
| **Insecta** | 1 |
| **Orthoptera** | 2–4 |
| **Hymenoptera** | 5–8 |
| **Hemiptera** | 9–12 |
| **Coleoptera** | 13,14 |
| **Diptera** | 15–20 |
| **Lepidoptera** | 21–26 |

**Supplementary Table S3.** Transgenic strains generated in this study.

| Line | Substitution(s) | Genotype | M/M viable? | Bang Sensitivity |  | Ouabain Sensitivity (Adult Exposure) |  |
| --- | --- | --- | --- | --- | --- | --- | --- |
|  |  |  |  | M/+ | M/M | M/+ | M/M |
| 0 | Δ2-6 | w<sup>118</sup>;Δ2-6/TM6 | no | yes | . | . | . |
| 1 | None | w<sup>118</sup>;GE-WT | yes | . | ref | . | ref |
| 20 | N122H | w<sup>118</sup>;N122H/TM6 | no | yes | . | +++ | . |
| 21 | Q111L | w<sup>118</sup>;Q111L/TM6 | no | yes | . | +++ | . |
| 22 | Q111T | w<sup>118</sup>;Q111T/TM6 | no | yes | . | . | . |
| 23 | Q111V | w<sup>118</sup>;Q111V/TM6 | no | yes | . | +++ | . |
| 24 | C104Y | w<sup>118</sup>;C104Y/TM6 | no | yes | . | . | . |
| 25 | Q111T,N122H | w<sup>118</sup>;Q111T, N122H/TM6 | no | yes | . | . | . |
| 26 | Q111V,N122H | w<sup>118</sup>; Q111V, N122H/TM6 | no | yes | . | . | . |
| 27 | C104Y,N122Y | w<sup>118</sup>;C104Y,N122Y/TM6 | no | yes | . | . | . |
| 28 | N122Y | w<sup>118</sup>;N122Y/TM6 | no | yes | . | . | . |
| 31 | A119S | w<sup>118</sup>;A119S | yes | no | yes | ++ | + |
| 32 | Q111L,A119S | w<sup>118</sup>;Q111L,A119S | yes | no | yes | . | + |
| 33 | Q111V,A119S | w<sup>118</sup>;Q111V,A119S | yes | no | yes | + | ++ |
| 34 | A119S,N122H | w<sup>118</sup>;A119S,N122H | yes | no | yes | + | + |
| 35 | Q111T,A119S | w<sup>118</sup>; Q111T,A119S | yes | no | yes | . | + |

NOTE – “TM6” = *TM6B, P{w[+mC]=Dfd-EYFP}3, Sb<sup>1</sup>,Tb<sup>1</sup>,ca<sup>1</sup>*. Δ2-6 is a loss-of-function deletion. M/+ and M/M refer to heterozygous and homozygous substitutions, respectively. Mut/+ were assayed such that the + allele is derived from line 1 (the engineered wild-type strain). “ref” refers to the reference strain used for the assay. “.” indicates genotypes that were not assayed. Sensitivity of adults to ouabain exposure is qualitatively scored on a scale from highest = “+++” to lowest = “+”.

